# Inter-organellar nucleic acid communication by a mitochondrial tRNA regulates nuclear metabolic transcription

**DOI:** 10.1101/2023.09.21.558912

**Authors:** Christopher Rouya, King Faisal Yambire, Mark L. Derbyshire, Hanan Alwaseem, Sohail F. Tavazoie

## Abstract

Efficient communication between mitochondria and the nucleus underlies homoeostatic metabolic control, though the involved mitochondrial factors and their mechanisms are poorly defined. Here, we report the surprising detection of multiple mitochondrial-derived transfer RNAs (mito-tRNAs) within the nuclei of human cells. Focused studies of nuclear-transported mito-tRNA-asparagine (mtAsn) revealed that its cognate charging enzyme (NARS2) is also present in the nucleus. MtAsn promoted interaction of NARS2 with histone deacetylase 2 (HDAC2), and repressed HDAC2 association with specific chromatin loci. Perturbation of this axis using antisense oligonucleotides promoted nucleotide biogenesis and enhanced breast cancer growth, and RNA and nascent transcript sequencing demonstrated specific alterations in the transcription of nuclear genes. These findings uncover nucleic-acid mediated communication between two organelles and the existence of a machinery for nuclear gene regulation by a mito-tRNA that restricts tumor growth through metabolic control.

**Highlights:**

1. Multiple mitochondrial-derived tRNAs are detected in human cell nuclei
2. MtAsn promotes binding between NARS2 and HDAC2
3. Metabolic alterations driven by mtAsn impact cell proliferation
4. MtAsn inhibition releases HDAC2 to bind and transcriptionally regulate multiple nuclear genes

## INTRODUCTION

Signaling between organelles coordinates cellular responses to nutrient imbalance, inflammatory insults, and various other stressors, and communication between the mitochondria and nucleus harmonizes mitochondrial metabolism with nuclear gene expression control ^1–4^. The human mitochondrial genome encodes for 13 essential components of the electron transport chain (ETC) whose expression must be coordinated with the majority of subunits encoded by the nuclear genome ^3, 5, 6^. In addition, two ribosomal RNAs (rRNAs) and 22 mito-tRNAs are expressed by the mitochondrial genome, contributing to the mitochondrial-specific translation apparatus ^5^. Nuclear control of mitochondrial biogenesis and metabolic activity is accomplished through so-called anterograde regulation, where transcription factors such as TFAM, NRF1, and peroxisome proliferator-activated receptors (PPARs) orchestrate the expression of mitochondrial respiratory chain genes, mitochondrial protein import and assembly factors, and components of the mitochondrial transcription and translation machineries ^7, 8^. Reciprocal control is achieved through the retrograde response, where mitochondrial-derived signaling factors initiate cytoplasmic or nuclear mechanisms of gene expression regulation to ultimately feedback on mitochondrial function ^7^.

While examples of retrograde signaling are well-described in lower eukaryotes, mammalian mito-nuclear factors and their mechanisms of action are still emerging. In particular, mitochondrial nucleic acids have become increasingly appreciated as mediators of mito-nuclear control. Release of mitochondrial DNA into the cytosol can engage the cGAS-STING pathway and drive transcriptional networks to support inflammatory processes ^9, 10^. Mitochondrial membrane translocation of nucleic acids depends upon outer mitochondrial membrane pore structures such as the oligomeric BAX/BAK and VDAC1 channels, and can involve the herniation of regions of the inner mitochondrial membrane harboring mitochondrial DNA ^11–14^. Several studies have also observed mitochondrial-derived RNAs present within the cytoplasm. In flies and human cells, impaired clearance of mitochondrial double-stranded RNAs (dsRNA) results in their accumulation and release via BAX/BAK pores into the cytosol, where binding to the MDA-5 RNA sensor triggers inflammatory signaling ^15, 16^. Sequencing of RNAs bound by the cytosolic dsRNA-sensing kinase, PKR, revealed pervasive interactions with mitochondrial dsRNA which contributed to PKR-dependent stress signaling and the development of neuroinflammation in a model of huntington’s disease ^17, 18^. Efflux of mitochondrial RNA into the cytoplasm is also associated with responses to genotoxic stressors, chondrocyte function, and thermogenesis in adipocytes ^19–21^. Non-coding mitochondrial RNAs can also reach the nucleus, although their mechanistic impact on nuclear gene expression control and functional relevance remain largely ill-defined ^22, 23^.

TRNAs are small RNAs that decode mRNAs during translation and are enzymatically coupled to amino acids by aminoacyl tRNA synthetase (ARS) enzymes ^24^. Distinct sets of tRNAs are transcribed from the nuclear and mitochondrial genomes and undergo maturation through biogenesis pathways that involve compartment-specific factors. The prevailing model for tRNA compartmentalization suggests that mature tRNAs are present only in translationally active regions of the cell, and are excluded from the nucleus. In challenge of this view, work in Saccharomyces cerevisiae has observed bidirectional, nutrient-responsive mature tRNA transport between the cytoplasm and nucleus ^25–30^. In addition, cytosolic tRNAs can translocate to the mitochondria in organisms that lack some or all of their mito-tRNA genes, to maintain normal mitochondrial protein synthesis ^31–33^. Moreover, the threonine and methionine mito-tRNAs associate with RNA-binding proteins in the cytosol of human cells ^34^. Despite reported instances of multidirectional tRNA transport, an understanding of global tRNA localization patterns has remained elusive due to challenges in applying high-throughput methodologies to measure tRNA abundances. We have herein applied splinter ligation-based hybridization and tRNA-sequencing ^35^ to subcellular fractions in order to catalogue the distribution of tRNAs within cellular compartments. This has uncovered mitochondrial tRNA localization within the human cell nucleus. We observe the abundant nuclear localization of mtAsn along with its charging enzyme, NARS2, and find that this mito-tRNA promotes interaction between NARS2 and the epigenetic regulator, HDAC2. Notably, antisense oligonucleotide-mediated inhibition of the NARS2/mtAsn/HDAC2 axis establishes a metabolic program that supports enhanced breast cancer growth. Moreover, inhibition of mtAsn demonstrates a role for this tRNA in eviction of HDAC2 from target gene loci to alter nuclear transcription.

## RESULTS

### Mitochondrial-derived tRNAs localize to human cell nuclei

We performed tRNA profiling of small RNA populations enriched from the cytoplasmic and nuclear fractions of the MCF10A breast epithelial cell line, the MDA-MB-231 breast cancer cell line and the *in vivo* selected, highly metastatic isogenic MDA-LM2 sub-line. Multiple mito-tRNAs were detected at high abundance in the nuclei of all cells analyzed, with the mito-tRNAs for asparagine (Asn), glutamic acid (Glu), and serine (Ser) observed most prominently within nuclear fractions (Figs. 1A-1B, S1A-1B). Consistent with these tRNA profiling results, mito-tRNAs Asn, Glu, and Ser_TGA_ were also detected in nuclear fractions by Northern blot analysis, while mito-tRNAs encoding for lysine and tryptophan were not (Figs. 1C, S1C). To validate these findings using a third orthogonal approach, we adapted established approaches to detect tRNA using RNA fluorescence in situ hybridization (RNA-FISH) ^25, 36–38^. Using locked nucleic acid probes to enhance specificity of signal ^39, 40^, we performed RNA-FISH to measure mtAsn localization, and confirmed the presence of mtAsn within nuclei (Fig. S1D). Conversely, mtLys was restricted to the cytoplasm and mitochondria, consistent with the aforementioned fractionation studies (Fig. S1D). Nuclear-encoded tRNA Asn_GTT_ served as a control—being enriched in the cytoplasm and detectable in the nucleus (Fig. S1D). Thus, an unbiased profiling strategy coupled with orthogonal validation approaches identified nuclear-localized mito-tRNAs in multiple human cell lines.

**Figure 1.**
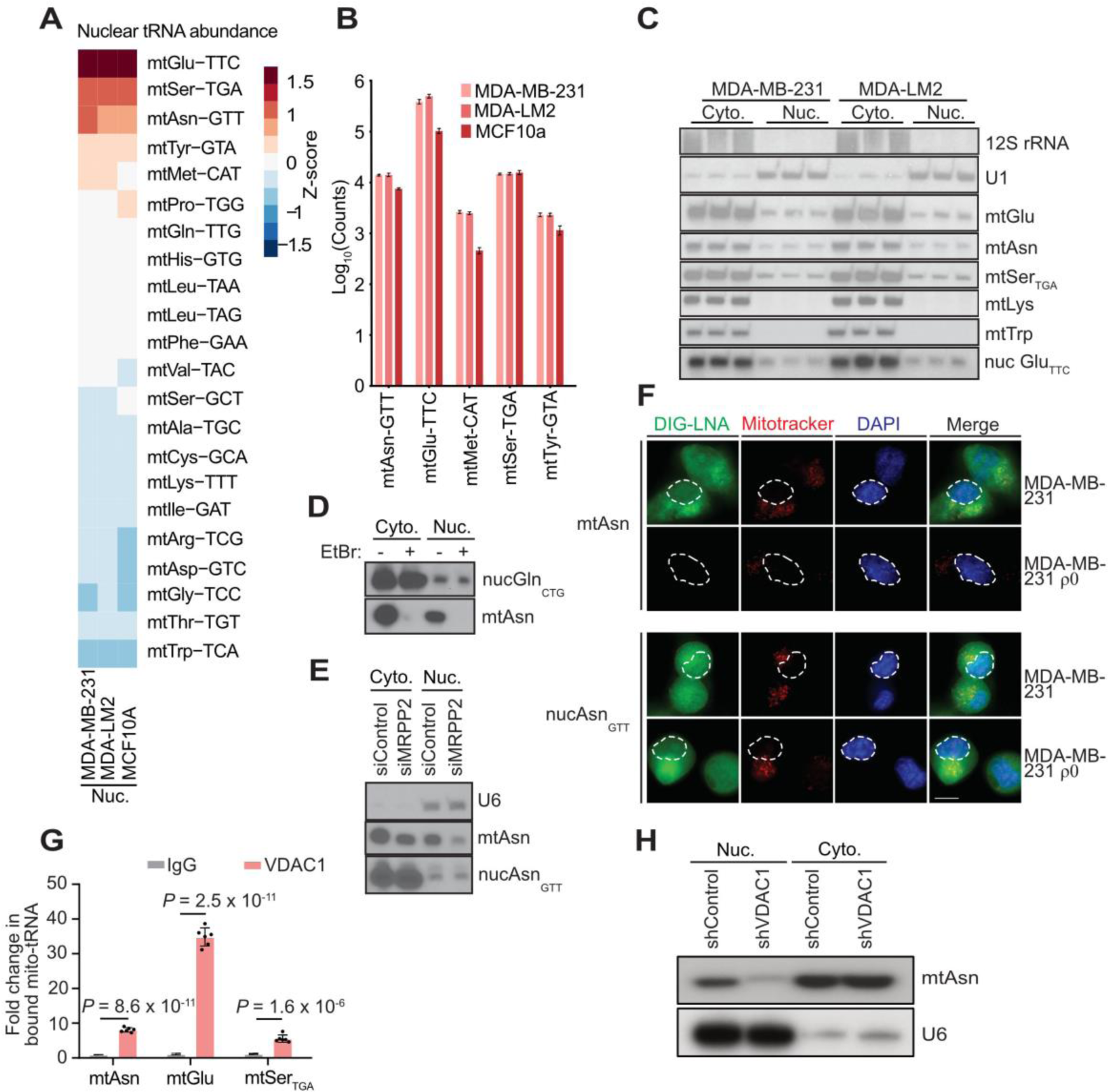
Mitochondrial translation components localize to cell nuclei. A. Mito-tRNA profiling of nuclear fractions for indicated cell lines. B. Average nuclear count data for mito-tRNAs highly abundant within nuclear fractions. C. Northern blot of cytoplasmic/mitochondrial and nuclear fractions to detect abundance of indicated RNAs. mt = mitochondrial-encoded, nuc = nuclear-encoded D. Northern blot analysis of cytoplasmic/mitochondrial and nuclear fractions upon treatment with ethidium bromide (200 ng/mL) for 48 hours in MDA-MB-231 cells. E. HEK293 cells transfected with a control siRNA or an siRNA targeting MRPP2, were fractionated into cytoplasm, nuclei, and mitochondria. RNA isolated from each fraction was subjected to Northern blot analysis for the indicated RNAs. F. RNA FISH of mtAsn and nuclear-encoded tRNA Asn_GTT_ localization in MDA-MB-231 and MDA-MB-231 ρ0 cells. DAPI and Mitotracker visualize the nuclei and mitochondria of cells, respectively. Dotted outlines highlight nuclear position, scale bar = 10 μm. G. YAMAT quantitative PCR (YAMAT-qPCR) was performed for the indicated mito-tRNAs from IgG or VDAC1 immunoprecipitates, normalized to total tRNA abundance. P-values are derived from two-sided Student’s t-test, *n* = 6. H. Total RNA from nuclear or cytosolic/mitochondrial fractions of cells transduced with small hairpin RNAs (shRNAs) against VDAC1 or a non-targeting control was subjected to Northern blotting.

Inhibition of mitochondrial transcription using low doses of ethidium bromide (EtBr;^41^) resulted in complete loss of mito-tRNA expression within the nuclei of EtBr-treated cells (Fig. 1D). Moreover, disrupting mito-tRNA processing through siRNA- or shRNA-mediated depletion of a component of the mitochondrial tRNA processing machinery, MRPP2, elicited specific and robust reductions in mito-tRNA levels in nuclei (Figs. 1E, S2A-2C) ^42^. RNA-FISH carried out with MDA-MB-231 ρ0 cells lacking transcriptionally active mitochondria (Fig. S2D) also abolished mtAsn expression in both the cytoplasmic and nuclear compartments, while nuclear-encoded tRNA Asn_GTT_ levels were unchanged (Fig. 1F). Thus, suppression of mitochondrial transcription or tRNA processing using multiple orthogonal approaches reduced the abundance of nuclear-localized mito-tRNAs, supporting a model whereby transcription and transport of specific mito-tRNAs from the mitochondria to the nuclei of human cells determines nuclear mito-tRNA abundance.

Escape of small mitochondrial DNA fragments into the cytoplasm can occur via translocation through VDAC1 oligomeric channels in the outer mitochondrial membrane ^12^. Given that a positively charged, N-terminal region of VDAC1 monomers forms a channel that permits flow of ∼100 nucleotide-long mito-DNA fragments into the cytosol, we reasoned that negatively charged tRNA molecules of similar length may also co-opt this transport machinery. Consistent with this, mito-tRNAs were detected in VDAC1 immunoprecipitates, indicating that the VDAC1 pore can indeed associate with mito-tRNAs (Fig. 1G). Moreover, we observed a substantial impairment of nuclear-localization for mito-tRNAs following shRNA-mediated depletion of the VDAC1 channel and upon treatment with the VDAC1 inhibitor, VBIT-4 (Figs. 1H, S2E-2F)^43^. These findings implicate the mitochondrial VDAC1 pore in facilitating mito-tRNA transport from the mitochondria to the nucleus.

### A NARS2-HDAC2 complex depends upon mtAsn

Mito-tRNAs undergo all processing and modification steps in the mitochondria and are eliminated by the organellar RNA decay machinery, suggesting that nuclear localization is not a requirement for mito-tRNA biogenesis or destabilization ^44^. We thus hypothesized that within the nucleus, mito-tRNAs may participate in gene regulation by interacting with specific effector proteins. We conducted focused mechanistic studies on mtAsn, a highly nuclear abundant mito-tRNA (Figs. 1A-1B). We reasoned that the ideal mito-tRNA-interacting partner would belong to the class of proteins that has evolved to recognize mito-tRNAs with high specificity: mitochondrial aminoacyl tRNA synthetases (ARS2). We generated nuclear and cytoplasmic fractions of MDA-MB-231 cells and immunoblotted for NARS2, the aminoacyl tRNA synthetase for mtAsn. As expected, NARS2 was detected in cytoplasmic fractions, which contain mitochondria (Fig. 2A). Remarkably, we also detected NARS2 in nuclear fractions, an observation corroborated by immunofluorescence experiments (Figs. 2A-2B). Analysis of publicly available spatial imaging datasets corroborated NAARS2 nuclear localization^45^. These findings reveal that a mitochondrial tRNA charging enzyme is present within the nuclei of human cells, where its cognate mito-tRNA also resides.

**Figure 2.**
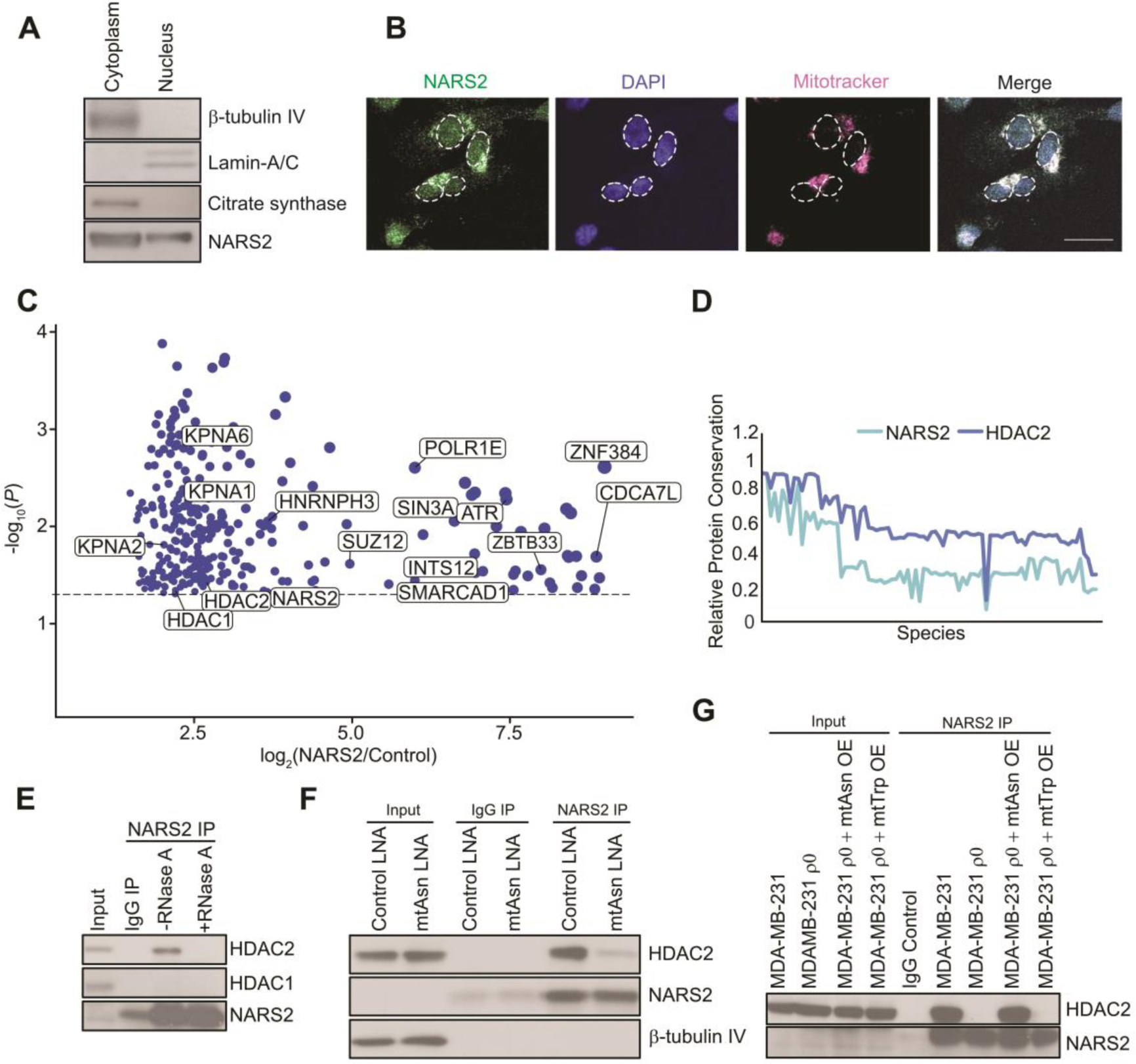
Mitochondrial asparaginyl-tRNA synthetase NARS2 forms a nuclear network of protein interactions. A. Cytoplasmic/mitochondrial and nuclear lysates from MDA-MB-231 cells were assessed via immunoblotting for the labeled proteins. B. Immunofluorescence in MDA-MB-231 cells for the indicated factors. Scale bar = 20 μm. C. Plot of high confidence nuclear-localized, interacting partners identified using APEX2-NARS2-mediated biotinylation experiments. Proteins of interest with roles in nucleocytoplasmic trafficking or transcriptional regulation are labelled. D. Phylogenetic profile of NARS2. Relative protein sequence conservation to humans is shown for NARS2 (light blue) and the co-evolved HDAC2 (dark blue). X-axis depicts the ∼80 species/genomes compared (species names omitted from plot due to space, see ^49^). E. NARS2 immunoprecipitation was performed, and immunoblotting carried out for the labeled factors. Immunoprecipitation reactions were treated with or without RNase A overnight. F. MDA-MB-231 lysates were derived from cells transfected with control or mito-tRNA Asn-targeting LNA. Lysates were subjected to NARS2 immunoprecipitation and immunoblotting for the indicated factors. G. NARS2 immunoprecipitations were performed in MDA-MB-231 or MDA-MB-231 ρ0 cells transfected with control vector, mito-tRNA Asn, or mito-tRNA Trp over-expression vectors. Immunoblotting was performed for the indicated proteins.

We next sought to identify proteins that interact with NARS2 that might transport it or mediate the nuclear effects of a mtAsn-NARS2 complex. To this end, we employed APEX2 technology ^46^, and expressed a NARS2-APEX2 fusion construct in HEK293 cells followed by rapid biotin proximity labelling of NARS2-associated proteins, streptavidin pulldown, and mass spectrometry. In parallel, we carried out experiments using mitochondrial targeting signal (MTS) and nuclear localization signal (NLS) APEX2 fusion proteins to fully catalogue proteins in each sub-compartment. Our APEX2 data represent the first analysis of the NARS2-associated proteome, and approximately 480 high confidence NARS2-associated proteins (fold change > 3, p-value<0.05) were specific to the NARS2-APEX2 construct with most of these factors demonstrating mitochondrial localization, as expected (Figs. S3A-3D). Notably, a subset of NARS2-binding proteins exhibited strong nucleoplasmic localization based on overlap with our NLS-APEX2 data, as well as known GO annotations for protein localization (Fig. 2C). Cytosolic-nuclear transport can be mediated by an importin-α/importin-β complex in association with NLS-harboring cargoes ^47^. Our mass spectrometry data identified NARS2 binding to several importin-α family members, and through co-immunoprecipitation we validated an interaction with the importin-α subunit factor KPNA2 (Fig. S3E), suggesting that transport of NARS2 to the nucleus may be mediated by the importin-α/β nuclear import machinery.

Groups of proteins that function in common molecular complexes or pathways have been shown to exhibit co-variation in protein sequence conservation when surveyed across phylogenetic clades ^48, 49^. We performed such a co-evolution analysis on NARS2 using the Phylogene platform ^49^. Intriguingly, the second most highly co-evolved gene with NARS2 across all genes was histone deacetylase 2 (HDAC2) (Pearson correlation 0.98; Fig. 2D, Table S1), a nuclear factor that was also detected in our aforementioned NARS2-APEX2 dataset (Fig. 2C). APEX2 technology allows for a more complete catalog of bound proteins, including weak or transient interactions, perhaps explaining why previous HDAC2 affinity-mass spectrometry-based methods failed to detect NARS2 ^50^. HDAC2 can alter chromatin compaction through epigenomic changes in histone modification, resulting in downstream transcriptional repression upon the removal of histone acetylation groups. Genomic occupancy studies of histone deacetylases such as HDAC2 report binding to transcriptionally active genes as well, which suggests that the recycling of histone acetylation marks may be necessary to augment transcription ^51–54^. Co-immunoprecipitations (co-IP) of either endogenous NARS2 or HDAC2 revealed a specific NARS2-HDAC2 interaction, which was RNA dependent (Figs. 2E, S4A). Co-localization and binding could be observed in both the nucleus and cytoplasm via fractionation followed by co-IP and proximity ligation assay, respectively (Figs. S4B-4C). *In vitro* co-precipitation experiments with recombinant SUMO-tagged NARS2 and HDAC2 purified from *E. coli* and insect cells failed to detect direct contact between the two proteins, suggesting requirement of additional factor(s) for interaction (Fig. S4D). Strikingly, the NARS2-HDAC2 interaction was abrogated in ρ0 cells, suggesting that a mitochondrial nucleic acid is, at least in part, responsible for this interaction (Fig. S4E). We hypothesized that mtAsn may contribute to complex formation, and to this end transfected cells with a Locked Nucleic Acid (LNA) targeting mtAsn to interfere with its binding to NARS2-HDAC2. The inability of exogenous nucleic acids such as LNAs to enter mitochondria ^55^ provided a means for exclusively targeting mito-tRNAs outside of the mitochondria. Indeed, LNA inhibition did not impair the translation of mitochondrial-encoded COX II or nuclear-encoded OXPHOS subunits whose expression and assembly depends upon efficient mitochondrial translation (Fig. S4F) ^3^. These data suggest that LNA transfection did not interfere with the canonical mitochondrial function of the targeted mito-tRNAs. While cells transfected with a control, non-targeting LNA maintained the NARS2-HDAC2 interaction, LNA-mediated inhibition of mtAsn reduced complex formation (Fig. 2F). To assess whether this mito-tRNA was sufficient to mediate association between NARS2 and HDAC2, we restored expression of mtAsn in ρ0 cells via transient transfection using a previously described system (Fig. S4G) ^56^. Notably, sole expression of mtAsn was sufficient to rescue the NARS2-HDAC2 interaction in ρ0 cells, while expression of the control mtTrp failed to rescue the interaction (Fig. 2G), consistent with a model whereby NARS2 together with its associated mito-tRNA interacts with HDAC2. These results reveal that NARS2 interacts with HDAC2 in both the nucleus and cytoplasm of human cells, and that binding is driven by mtAsn.

### MtAsn inhibits cell proliferation

Intriguingly, we noted that mtAsn inhibition stimulated proliferation in multiple cancer cell lines (Figs. 3A, S5A-5B). LNAs are highly potent binders of RNAs and can establish long-lasting and specific depletion of RNAs. Consistent with this, a single transfection of mtAsn-targeting LNA augmented *in vitro* proliferation for up to 20 days (Fig. 3B). Conversely, transient overexpression of mtAsn suppressed cell proliferation *in vitro* (Fig. S5C). Proliferative enhancement upon mtAsn inhibition was abrogated by HDAC2 depletion, further supporting an epistatic interaction between HDAC2 and mtAsn (Fig. S5D-5E).

**Figure 3.**
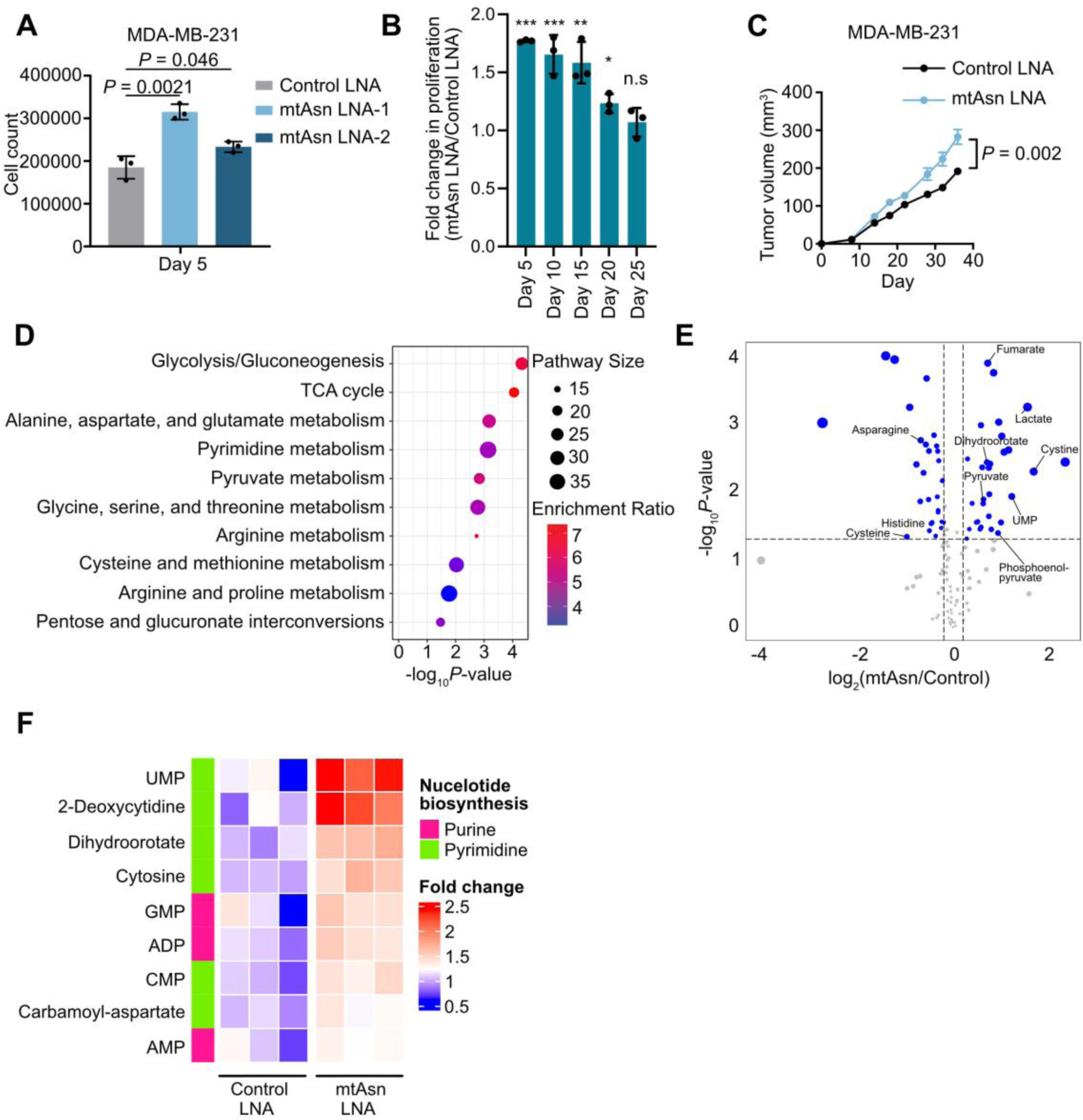
mtAsn inhibition promotes cancer cell proliferation. A. *In vitro* proliferation assay upon mtAsn inhibition. *n* = 3, p-values calculated using two-sided Student’s t-test. B. MDA-MB-231 cells were transfected with control or mtAsn LNAs once, and split regularly. On days 2, 7, 12, 17, and 22, 50,000 cells were seeded into 6-well plates, and cells were subsequently counted on days 5, 10, 15, 20, 25. *n* = 3, p-values determined by two-sided Student’s t-test. C. 50,000 MDA-MB-231 cells transfected with indicated LNAs were transplanted into the mammary fat pads of mice, and primary tumor growth measured. n = 7 per group, p-value calculated by two-sided Student’s t-test. D. Metabolic pathway analysis for enriched metabolites in MDA-MB-231 cells transfected with mito-tRNA Asn LNA as compared to control LNA. E. Volcano plot depicted significantly altered metabolites upon mito-tRNA Asn inhibition. Blue-coloured circles indicate statistically significant metabolite alterations. F. Heat map representation of fold changes in total metabolite abundance for the indicated nucleotides and nucleotide intermediates in MDA-MB-231 cells transfected with non-targeting control or mtAsn-targeting LNAs.

Previous work has described widespread depletion of mitochondrial DNA copy number and transcripts across a variety of tumor types that is of unclear functional significance ^57–59^. We thus tested whether inhibiting mtAsn could impact tumor growth *in vivo*. In accordance with our *in vitro* findings, inhibition of mtAsn enhanced *in vivo* mammary tumor growth of MDA-MB-231 and HCC-1806-LM2c cells (Figs. 3C, S5F). These data reveal that perturbing a mito-tRNA nuclear signaling axis can induce a proliferative response.

### Metabolic regulation by mtAsn impacts proliferation

Bidirectional signaling between mitochondria and the nucleus maintains metabolic balance under several environmental perturbations^60–70^, suggesting that differential metabolite usage may underlie the *in vitro* and *in vivo* proliferative advantage of mtAsn-inhibited cells. We thus profiled total metabolites in MDA-MB-231 cells upon mtAsn inhibition. Unsupervised hierarchical clustering revealed mtAsn-dependent metabolic modulation in MDA-MB-231 cells (Fig. S6A). Inhibition of mtAsn modulated the levels of metabolites enriched in glycolysis, TCA cycle, amino acid metabolism and pyrimidine biosynthesis— key pathways with established growth promoting effects (Figs. 3D-3F). Consistent with induced glycolytic and TCA cycle activities, mtAsn inhibition increased the levels of several glycolytic and TCA cycle metabolites (Figs. S6B, 3D). Accordingly, suppressing mtAsn-dependent retrograde signaling enhanced basal, maximal, and ATP-coupled oxygen consumption (mitochondrial respiration) and extracellular acidification (glycolytic) rates of MDA-MB-231 cells (Figs. S6C-6D). Thus, perturbing the mtAsn mito-nuclear axis can significantly drive mitochondrial respiration and impact cellular metabolic physiology.

### The NARS2/mtAsn/HDAC2 axis controls nuclear gene transcription

To understand how mtAsn may mechanistically control metabolism through the tRNA-dependent NARS2-HDAC2 interaction, we carried out transcriptomic profiling on breast cancer cells transfected with either a non-targeting LNA, two independent LNAs complementary to the 5’ or 3’ ends of mtAsn for inhibiting the tRNA, and an LNA that interacts with the non-nuclear-localized mtLys (Figs. S7A-7C, 1C). Consistent with direct targeting, we observed a highly significant overlap of differentially expressed genes between the mtAsn LNA-1 and LNA-2 transcriptomes (Spearman correlation coefficient = 0.92, p = 1.05×10^-241^; Fig. S7E). Using mtLys-regulated genes as a control filter, we identified 574 genes that were specifically altered by mtAsn inhibition (Figs. S7E-7G). Amongst induced genes within this set, GO pathways enriched included mitotic cell cycle and nuclear division, indicating a potential role for mtAsn in regulating proliferation (Fig. S7H). We performed an additional RNA-seq analysis of cells depleted of NARS2 by RNAi (Figs. S7I-7K), reasoning that the genes downstream of the NARS2-mtAsn-HDAC2 complex would be similarly regulated by both transient NARS2 depletion and mtAsn inhibition. Analysis of these RNA-seq datasets uncovered ∼130 genes modulated by both NARS2 and mtAsn (Fig. S7L). These findings reveal a regulon of nuclear-encoded genes controlled by a mitochondrial tRNA and its aminoacyl tRNA synthetase.

To determine whether a NARS2-tRNA-HDAC2 complex might alter gene expression via regulated association with chromatin, we sought to define the genomic localization of the transcriptional regulator HDAC2 upon mito-tRNA inhibition using CUT&RUN ^71^. Inhibition of mtAsn caused a global enhancement of HDAC2 occupancy near transcriptional start sites (Figs. 4A, S8A). Consistent with these findings, LNA-mediated mtAsn inhibition enhanced HDAC2 abundance in chromatin fractions (Fig. S8B). Amongst the ∼130 genes regulated by mtAsn and NARS2, ∼60 exhibited altered HDAC2 binding at their promoter regions and GO enrichment analysis of HDAC2-bound targets identified mtAsn- and NARS2-dependent regulation of genes associated with glutamate biosynthesis (Fig. 4B), a pathway upstream of the aforementioned observed metabolic rewiring (Figs. 3D and S6B). We next performed nascent transcript profiling by global run-on sequencing (GRO-seq) to assess whether the mtAsn axis directly influenced nuclear transcription of these downstream targets. Pervasive changes in transcription upon inhibition of mtAsn were detected (Fig. 4C), and we noted several target transcripts whose abundances were regulated by NARS2/mtAsn at the mature mRNA level also exhibited changes in transcription rate (Fig. 4D). These data implicate mtAsn as a regulator of nuclear transcription and suggest that it may influence cellular proliferation and metabolism through changes in glutamate biosynthesis.

**Figure 4.**
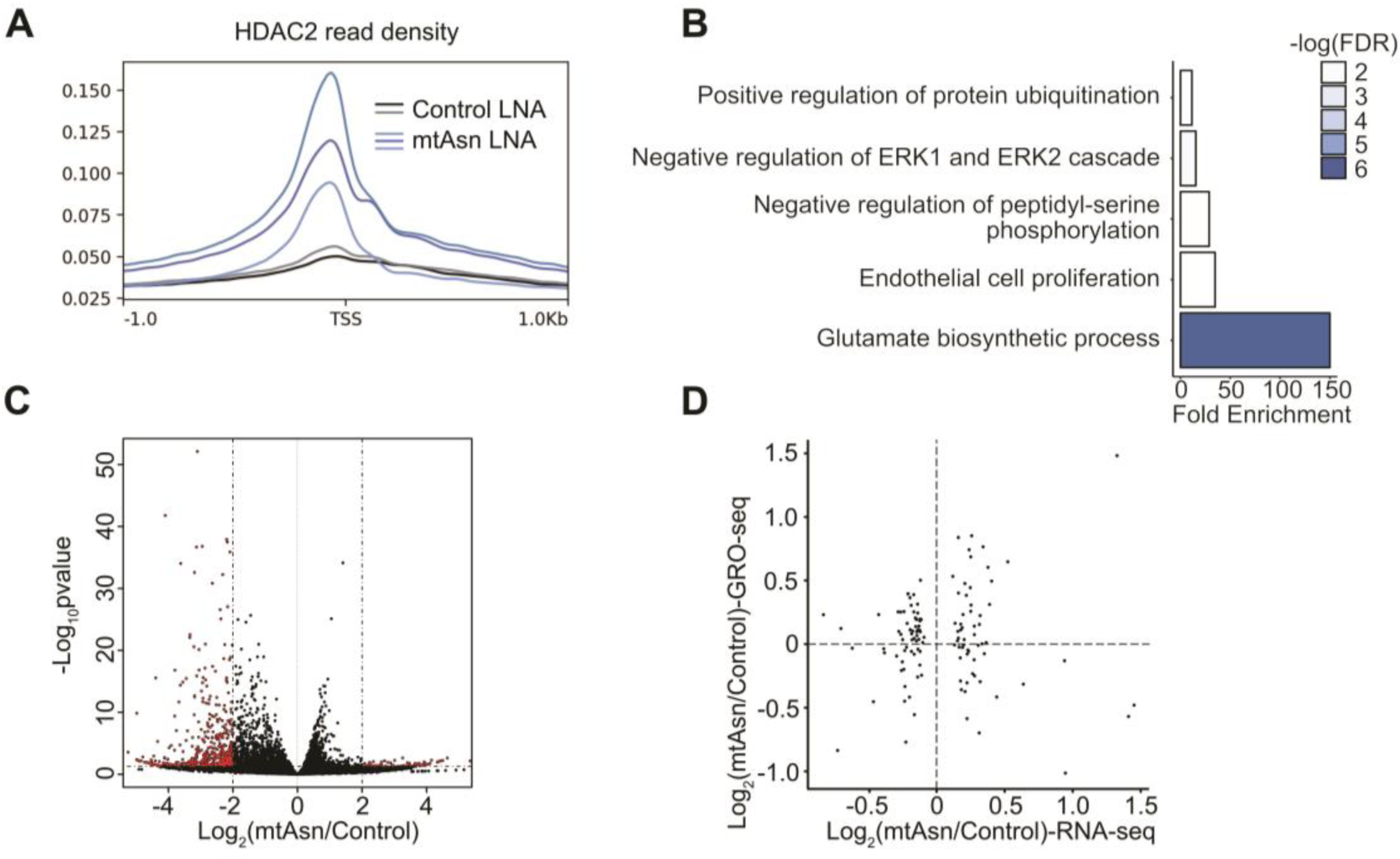
A mito-tRNA-dependent NARS2-HDAC2 complex drives a specific gene expression program via transcriptional control and suppresses growth. A. HDAC2 read density surrounding transcription start sites TSS) upon mtAsn inhibition. B. GO ontology of genes whose promoters are differentially bound by HDAC2 upon inhibition of mtAsn, as well as coordinately regulated at the mRNA level by mtAsn and NARS2. C. GRO-seq volcano plot depicting genes with significantly altered (red) normalized read counts. Dotted lines represent log2(foldchange) of −2 and 2, respectively. D. Scatter plot depicting GRO-seq and RNA-seq log2 fold changes for genes commonly altered by NARS2 depletion and mtAsn LNA transfection. Dotted lines denote log2 fold changes of 0 for both datasets.

## DISCUSSION

Nuclear sensing of mitochondrial status is critical for effective control of cellular metabolism. Our tRNA profiling strategy has uncovered the nuclear localization of several mitochondrial tRNAs, and supports an RNA-mediated mechanism that signals across two organelles to alter cellular metabolism and proliferation. Transport of mito-tRNAs to the nucleus to regulate transcription represents an additional mitonuclear signaling arm, in conjunction with factors such as metabolites and chromatin-localized, metabolic enzymes ^62, 68–70, 72, 73^. Our findings reveal how a complement of mitochondrial tRNAs can exit the mitochondria through VDAC1 pores and enter the nucleus. Mito-tRNA-dependent assembly of a NARS2-HDAC2 complex evicts the HDAC2 enzyme from target genes, supporting a gene expression program that may curb cancer cell proliferation. Employing metabolite profiling experiments, we demonstrate how this tRNA-dependent, mitonuclear pathway elicits profound changes in cellular metabolism and proliferative capacity, enhancing the abundance of glycolytic and TCA cycle intermediates, as well as nucleotide precursors. Thus, our data supports an RNA-mediated, biochemical mechanism that signals across two organelles to reshape cellular metabolism and proliferation.

The NARS2-mito-tRNA-HDAC2 interaction implicates a mitochondrial-derived RNA in chromatin-based gene regulation. Although HDAC2 is a transcriptional regulatory protein, it has been found to exhibit RNA binding capacity, which was reported to aid in HDAC2 engagement with chromatin and RNA polymerase II ^52, 74, 75^. Although the precise nature of HDAC2-associated RNA has not been defined, our work has identified a mitochondrial tRNA that regulates association of HDAC2 with chromatin to regulate the transcription of specific target genes. This HDAC2-dependent chromatin-based rewiring by mito-tRNA Asn additionally requires NARS2, but the complement of potential transcription factors that transmit a proliferative signal remains to be determined. Importantly, histone deacetylases have been previously shown to form nuclear-localized complexes with cytosolic tRNA synthetases ^76–78^. Cytosolic tRNA synthetases have evolved many additional domains that enable non-canonical binding to such factors to influence gene expression ^76–79^. In contrast, mitochondrial tRNA synthetases have not been described to have such additional domains that may mediate additional interactions, distinguishing this mitochondrial tRNA synthetase-based mechanism of transcriptional regulation from previously described cytosolic tRNA synthetase based processes. Further distinguishing is our finding that a tRNA governs this transcriptional regulatory response mediated by a cognate charging enzyme.

Our tRNA profiling strategy has uncovered the nuclear localization of several mitochondrial tRNAs. While VDAC1 oligomerization explains how these mito-tRNAs are released from mitochondria, how they might enter the nucleus remains unclear. In yeast, cytosolic tRNAs can re-enter the nucleus via the Transportin ortholog, Mtr10, suggesting a potential mechanism for mito-tRNA transport into human nuclei ^80^. Our detection of several mito-tRNAs and their cognate charging enzymes within the nucleus raises the intriguing possibility of additional mito-tRNA/charging enzyme axes regulating diverse nuclear processes in response to potential mitochondrial/metabolic perturbations. In many ways, these mito-tRNAs represent ideal mitonuclear signaling axes. Mitochondrial translation, which serves to generate important components of the oxidative phosphorylation machinery, depends on mito-tRNA abundance, while amino acid bioavailability influences mito-tRNA stability and function. Thus, mito-tRNAs can communicate a comprehensive picture of mitochondrial status, by signaling both mitochondrial translation capacity as well as the supply of specific amino acids within the mitochondria to the nucleus. Future work is needed to characterize the roles of other mito-tRNAs and their charging enzymes in mito-nuclear communication and control.

Depletion of mitochondrial transcripts has been observed across a wide range of tumor types, indicating that impairing mitochondrial RNA function can promote tumor growth in certain contexts. Through the transfection of LNA oligonucleotides which, as with most nucleic acids, do not appreciably enter the mitochondria, we find that extramitochondrial targeting of a single mito-tRNA can rewire cell metabolism and augment growth. TRNA abundance and aminoacylation could serve as surrogates for subcellular translation capacity and nutrient status, and in this way the mito-tRNA Asn axis described herein may allow for the integration of multiple mitochondrial cues to enact nuclear gene expression changes.

Our work supports a novel paradigm for transcriptional control by mito-tRNAs. In addition to roles in growth control and cancer, it is likely that a host of mitonuclear pathways, each governed by a unique mito-tRNA, underlie other physiological processes. Mutations in mito-tRNAs and their charging enzymes are associated with a spectrum of human disorders that affect the central nervous system and musculature ^81, 82^. While it has been postulated that defects in mitochondrial translation that arise from such specific mutations are the primary culprits for disease phenotypes, dysfunctional nuclear signaling and transcriptional deregulation may also contribute ^82^. For such disorders, as well as potentially other metabolic disorders with associated perturbations to this mitonuclear axis, the ability of oligonucleotides such as LNAs—as used herein to modulate mito-tRNA nuclear transcriptional control holds promise for previously unrecognized therapeutic applications. Discerning how other mitochondrial tRNAs or their charging enzymes might govern changes in nuclear gene expression and perhaps orchestrate additional cellular functions will be critical for providing a comprehensive understanding of inter-organellar communication and its dysregulation in disease.

## METHODS

### RESOURCE AVAILABILITY

#### Lead contact

Further information and requests for resources and reagents should be directed to and will be fulfilled by the lead contact, Sohail Tavazoie.

#### Materials availability

Plasmids generated in this study will be deposited to Addgene.

#### Data and Code Availability

All software is commercially available or cited in previous publications. Newly generated sequencing data has been deposited at GEO and are publicly available as of the date of publication. All other data will be available from the corresponding author upon request.

#### Cell Culture

HEK 293T, MDA-MB-231 and MDA-LM2 cell lines were cultured in DMEM supplemented with 10% FBS, glutamine, and pyruvate (D10F). MCF10A cells were cultured in DMEM/F12 media with the addition of 5% horse serum, 0.5 mg/ml of hydrocortisone, 100 ng/ml of cholera toxin, 10 µg/ml of insulin, and 20 ng/ml of EGF. HCC1806-LM2C cells were cultured in RPMI media containing 10% FBS, 1mM glucose, 10mM HEPES and 1mM sodium pyruvate. Cell lines were routinely tested for mycoplasma contamination and validated by STR analysis.

#### Mouse Experiments

All animal work was conducted in accordance with protocols approved by the Institutional Animal Care and Use Committee at The Rockefeller University. 5.0 × 10^4^ MDA-MB-231 or HCC1806-LM2C triple reporter-labeled cells were resuspended in a 1:1 mixture of PBS and Matrigel (BD Biosciences), and injected into the 4^th^ mammary fat pad of female NOD-SCID-g female mice 6-8 weeks of age. Tumor growth was measured twice weekly using digital calipers, with tumor volume calculated as (small diameter)^2^ × (large diameter) × Π / 6.

#### Generation of stable knockdown cell lines and LNA/siRNA transfection

For lentiviral-mediated gene knockdown, 5 µg of an shRNA vector (pLKO.1-puro vector-Sigma), 2 µg pMD2.g, and 3 µg psPAX2 were combined with 27 µL of lipofectamine 2000 (Thermo Fisher) in 200 µL of OPTI-MEM, and transfected into HEK 293T cells. 16 hours later, media was replaced, and 48 hours after transfection media containing lentiviral particles was collected and passed through a 0.45 µm filter. Virus-containing media was diluted 1:1 with fresh media, incubated with polybrene (10 µg/mL), and applied to cells plated in a 6-well dish at ∼25% confluency to begin infection. The day after infection, cells were cultured in D10F supplemented with 2 µg/mL of puromycin (Gibco) to begin selection. The following shRNAs were obtained from Sigma: HDAC2 (TRCN0000375947, TRCN0000039397), NARS2 (TRCN0000161788, TRCN0000163533, TRCN0000163609), MRPP2 (TRCN0000318935, TRCN0000318872), and VDAC1 (TRCN0000029126, TRCN0000029127).

500,000 MDA-MB-231 cells plated in a six-well dish were transfected with 20 nM of the appropriate siRNA (IDT) or 20 nM of appropriate LNA diluted in OPTI-MEM containing 7 µL of RNAiMAX (Thermo Fisher). Transfections were performed according to the manufacturer’s protocol. Two or three days following transfection, cells were harvested for downstream analyses.

#### Cell fractionation

To isolate cytoplasmic and nuclear fractions, cells were incubated in Buffer A (50 mM Tris-HCl pH 7.4, 2 mM MgCl2, 150 mM NaCl, 40 mM KCl, 10% glycerol, 0.1% TritonX-100, 0.1% Tween-40, and cOmplete protease inhibitor (Roche)) on ice for 10 minutes. Samples were briefly vortexed, and centrifuged at 1,300 x *g* for 10 minutes, and the supernatant was collected. This supernatant was centrifuged again at 10,000 x *g* and the resulting supernatant was kept as the cytoplasmic fraction. The nuclear pellet was washed twice with Buffer A, and then lysed with either RIPA buffer for protein analysis, or directly resuspended in Trizol reagent for RNA isolation.

Recovery of cytoplasm, mitochondria, and nuclear fractions was performed as previously described ^83^. Briefly, cells were resuspended in STM buffer (340 mM sucrose, 50 mM Tris-HCl pH 7.4, 2 mM MgCl2), and allowed to swell on ice for 15 minutes. Cytoplasmic membranes were mechanically disrupted by ∼30 strokes of a close-fitting Dounce homogenizer on ice, and the sample was spun at 1,300 x *g* for 10 minutes to pellet nuclei. The supernatant was centrifuged at 11,000 x *g,* and the cytoplasmic supernatant was collected. The pellet, containing the mitochondrial fraction, was washed twice with STM buffer and lysed in RIPA buffer or Trizol. The nuclear pellet was washed twice with STM buffer supplemented with 0.1% TritonX-100 and 0.5% Tween-40, and the nuclear pellet was lysed using RIPA buffer or Trizol.

To fractionate cytoplasm, nucleoplasm, and chromatin, cells were incubated in Buffer A on ice for 10 minutes and centrifuged at 1,300 x *g* for 10 minutes. The nuclear pellet was washed twice with Buffer A, and nucleoplasm was released by incubation in Buffer B (3 mM EDTA, 0.5 mM EGTA, 0.1 mM DTT) on ice for 30 minutes. Samples were centrifuged at 1,300 x *g* for 10 minutes, and the supernatant containing nucleoplasmic components was collected. The insoluble chromatin fraction was washed twice with Buffer B, and resuspended in 1X DNase I reaction buffer with 1 µL DNase I (NEB) for 15 minutes at room temperature, and inactivated with addition of 1 mM EDTA and 1% SDS.

#### Immunoprecipitation

Cells were incubated for 10 minutes on ice in cold lysis buffer (50 mM Tris-HCl pH 7.4, 150 mM NaCl, 1 mM EDTA, 0.5% NP-40, 10% glycerol, RNasein Plus (Promega), and cOmplete protease inhibitor), and briefly sonicated on low settings to enhance nuclear lysis. Lysate was centrifuged at 10,000 x *g* for 10 minutes, and 2 mg of supernatant was combined with 3 µg of target antibody or IgG control and rotated overnight at 4 degrees. For RNase A treatments, RNase inhibitor was omitted from reactions and 10 µg of RNase A (Thermo Fisher) was included. Subsequently, 30 µL of washed Protein G Dynabeads (Thermo Fisher) were added to each sample and incubated for 1 hour at 4 degrees with rotation. Beads were washed 3 times with lysis buffer, and resuspended in 1X LDS sample buffer with 1X reducing agent (Thermo Fisher), boiled for 8 minutes at 95 degrees to elute. For VDAC1 immunoprecipitation and subsequent YAMAT-RT-qPCR, cell pellets were pre-treated for 30 minutes with 2mM Ethylene glycolbis (succinimidylsuccinate) crosslinker prior to cell lysis as described above. RNA was then isolated from IgG control and VDAC1 immunoprecipitates on Protein G Dynabeads using TRIzol reagent.

#### RNA Isolation and measurement

For Northern analysis, RNA was isolated from cells using TRIzol or TRIzol LS reagents (Invitrogen). For qPCR analyses, RNA was isolated using the Total RNA Purification Kit (Norgen), and samples were DNase-treated on column following the manufacturer’s recommendations. Reverse transcription reactions were performed using SuperScript III (Life Technologies) with 200 ng–1 μg of RNA. cDNA was subjected to qPCR in 384-well plates (Axygen) in technical quadruplicate using FastSYBR Green Master Mix (Applied Biosystems) following the manufacturer’s instructions on a QuantStudio 5 Real-Time PCR System (Applied Biosystems). PrimeTime gene specific primers obtained from IDT were used in this study, and catalog numbers can be found in the supplementary methods. All qPCR experiments were performed with at least 3 biological replicates per condition. Measured data was analyzed using the ΔΔ*C*_T_ method. For RNA-seq, libraries were prepared using the TruSeq RNA Library Preparation Kit v2 (Illumina) according to the manufacturer’s instructions, and sequenced using the Illumina Nextseq 500 platform.

#### Proliferation assays

For proliferation experiments, three orthogonal approaches were used. 2.0 × 10^4^ of the indicated cells were seeded in triplicate in a 6-well plate. Viable cell numbers were then counted and quantified on day 3 or day 5 post-seeding using trypan blue and a hemacytometer. Alternatively, 3–5 x10^3^ cells were seeded in 96-well plates and viable cell numbers were determined using the Incucyte (Sartorius) Adherent-Cell-by-Cell module. These proliferation assays were complemented with the Cell Titer Glo (Promega) cell viability assay following modified manufacturer’s protocols. Briefly, 3–5 x10^3^ indicated cells were seeded in 96-well plates in 150μl of growth medium. On the day of seeding and at indicated timepoints, seeded cells were treated with 50μl of Cell Titer Glo reagent and luminescence was measured in a microplate reader. Fold increase in proliferation was determined as day 0-normalized luminescence at indicated timepoints.

#### CUT&RUN

CUT&RUN was performed as previously described (*28*). 0.5 million cells were resuspended in Wash Buffer (20 mM HEPES, pH 7.5, 150 NaCl, 0,5 mM Spermidine, cOmplete protease inhibitor) and incubated with 25 µL of concanavalin A-conjugated paramagnetic beads (Bangs laboratories) for 10 minutes at room temperature. Antibody Buffer (Wash buffer + 0.01% digitonin + 2 mM EDTA) containing mouse IgG or anti-HDAC2 antibody dilute 1:100 was then added to each sample, and overnight incubation at 4 degrees was carried out. Beads were then washed twice with Digitonin Buffer (Wash buffer + 0.01% digitonin), and then incubated with Digitonin Buffer containing 2.5 µL of CUTANA pAG-MNase (Epicypher) for 10 minutes at room temperature. Beads were subsequently washed twice with Digitonin Buffer and 1 µL of 100mM CaCl2 was added to initiate the cleavage reaction, performed for 2 hours at 4 degrees with rotation. Reactions were quenched using Stop Buffer (340 mM NaCl, 20 mM EDTA, 4 mM EGTA, 50 µg/mL RNase A, 50 µg/mL Glycogen), and cleaved DNA was purified using the ChIP Clean and Concentrator Kit (Zymo). Libraries were prepared with the NEBNext Ultra II Library Prep Kit for Illumina (NEB) and paired-end sequencing was performed using an Illumina Novaseq instrument.

#### ChIP-qPCR

MDA-MB-231 cells were fixed in 1% paraformaldehyde for 8 minutes, followed by quenching with glycine. Cells were then incubated on ice for 10 minutes in 1 mL of Lysis Buffer I (50 mM HEPES-KOH, pH 7.2, 140 mM NaCl, 1 mM EDTA, 10% glycerol, 0.5% NP-40, 0.25% TritonX-100) and centrifuged for 10 minutes at 500 x *g*. Cells were subsequently resuspended in 1 mL of Lysis Buffer 2 (10 mM Tris-HCl, pH 8, 200 mM NaCl, 1 mM EDTA, 0.5 mM EGTA) and incubated on ice for 10 minutes, followed by centrifugation at 1300 x *g* for 10 minutes at 4 degrees. Cells were then resuspended in 120 uL of Lysis Buffer 3 (10 mM Tris-HCl, pH 8.0, 100 mM NaCl, 1 mM EDTA, 0.5 mM EGTA, 0.1% Sodium deoxycholate, and 0.5% N-lauroylarcosine) and sonicated for a total of 5.5 minutes on ice (10 seconds on, 20 seconds off). ChIP-grade Protein G magnetic beads were washed twice with blocking solution (1 mg/mL BSA in 1X PBS with 0.05% Tween-20), and then incubated with IgG control or HDAC2 antibody at 4 degrees with rotation for 1 hour. Beads were washed again once with blocking solution. Chromatin fractions were centrifuged at max speed at 4 degrees for 10 minutes, and the supernatant was equally divided into Eppendorf tubes containing either beads bound to IgG or HDAC2 antibody. Incubations were performed overnight at 4 degrees, and the next day samples were washed 6 times with wash buffer (50 mM HEPES-KOH, 0.5M LiCl, 0.1% Sodium deoxycholate, 0.5% N-lauroylarcosine), once with TE + 50 mM NaCl (10 mM Tris-HCl pH 8.0, 1 mM EDTA, 50 mM NaCl), and then eluted twice with Elution buffer (50 mM Tris-HCl pH 8.0, 10 mM EDTA, 1% SDS). Samples were then treated with RNase A for 1 hour at 37 degrees, and proteinase K overnight at 65 degrees. DNA was subsequently purified using phenol:chloroform extraction, and subjected to qPCR reactions with gene-specific primers.

#### GRO-Seq

GRO-seq was performed as described previously ^84, 85^. Two 15-cm plates of 80% confluent MDA-MB-231 cells 2 days after transfection with non-targeting, mito-tRNA-Asn, or mito-tRNA-Lys LNAs per replicate were trypsonized and incubated on ice in swelling buffer (10 mM Tris-HCl, 2 mM MgCl2, 3 mM CaCl2) for 10 minutes on ice. Cells were centrifuged at 4 degrees for 10 minutes at 500 x *g,* and resuspended in 10 mL of lysis buffer (swelling buffer with 0.5% NP-40 and 10% glycerol) and incubated on ice for 10 minutes with gentle vortexing every two minutes. Centrifugation was performed at 4 degrees for 10 minutes at 1100 x *g,* and nuclei were washed once more with 10 mL of lysis buffer. After centrifugation, nuclei were resuspended in 1 mL of freezing buffer (50 mM Tris-HCl pH 8.3, 40% glycerol, 5 mM MgCl2, 0.1 mM EDTA), pelleted at 1100 x *g,* and resuspended once more in 100 uL of freezing buffer. Run-on was performed by addition of equal volumes of transcription buffer (10 mM Tris-HCl pH 8.0, 5 mM MgCl2, 1 mM DTT, 300 mM KCL, 20 units of SUPERase-In (Thermo Fisher), 1% Sarkosyl, 500 uM ATP, GTP, and Br-UTP, and 2 uM CTP), and incubated for 5 minutes at 30 degrees. RNA was extracted using Trizol, and subjected to base hydrolysis using NaOH. 1M Tris-HCl pH 6.8 was added to neutralize the reaction, and RNA was purified using p-30 RNase-free columns (BioRad), and treated with RQ1 RNase-free DNase (Promega), and purified again use p-30 columns. Antarctic phosphatase and 1 uL of SUPERase-In was then added to reactions, and treated at 37 degrees for 1 hour. Anti-BrdU agarose beads (Santa Cruz) were blocked using blocking buffer (0.5 X SSPE, 1 mM EDTA, 0.05% Tween-20, 1 mg/mL BSA, 0.1% PVP) for 1 hour at 4 degrees. Nascent RNA was added to beads and allowed to bind for 1 hour at 4 degrees with rotation. Beads were subsequently washed three times with wash buffer (0.5 x SSPE, 1 mM EDTA, 0.05% Tween-20, 150 mM NaCl), and twice in TET buffer (TE pH 7.4, 0.05% Tween-20). Nascent BrU-incorporated RNA was eluted with 4 x 125 uL elution buffer (20 mM DTT, 300 mM NaCl, 1 mM EDTA, 0.1% SDS, 5 mM Tris-HCl pH 7.5), and RNA was extracted using phenol:chloroform. RNA was then treated with T4 PNK for 1 hour at 37 degrees and precipitated again.

Poly-A tailing was performed using poly-A polymerase (NEB) for 30 minutes at 37 degrees, and reverse transcription was carried out using poly-T primer and Superscript III, according to the manufacturer’s instructions. Tailed RNA was combined with 1 uL dNTPs and 2.5 uL 12.5 uM poly-T primer, heated for 5 minutes at 65 degrees, and chilled on ice for 1 minute. 0.5 uL SUPERase-In, 3 uL 0.1 mM DTT, 2 uL 25 mM MgCl2, 2 uL 10X buffer, and 1 uL superscript III. RT was performed at 50 degrees for 1 hour, followed by incubation with 4 uL of exonuclease I at 37 degrees for 1 hour. Reaction products were gel purified, excised, and eluted overnight in elution buffer (TE buffer with 0.1% Tween-20 and 150 mM NaCl) and ethanol precipitated. cDNA was circularized using 0.5 uL CircLigase (Epicentre), 1 uL CircLigase buffer, 0.5 uL 1 mM ATP, 0,5 uL 50 mM MnCl2 for 1 hours at 60 degrees. Relinearization was performed using 1.5 uL of ApeI (NEB), and ssDNA was separated using TBE-urea gel electrophoresis and gel purified. ssDNA was then amplified using Phusion HiFi enzyme according to the manufacturer’s instructions, and DNA was sequenced using an Illumina NextSeq instrument.

#### Data analysis and visualization

Fastq file quality was assessed using FastQC, and reads were aligned to the hg38 human genome with STAR v.2.6.0a using default parameters. Read counts were obtained from mapped bam files using featureCounts .v.1.5.3, and DESeq2 was used to identify differentially expressed genes (DEGs). For CUT&RUN analysis, reads were trimmed using Trimmomatic v.0.39, and fastq files were aligned to the hg38 genome using Bowtie2, peaks were called using MACS2, and bam to bigwig conversion (normalizing by reads per bin) was carried out using deeptools. Differentially bound regions were identified using the Diffbind package. Heatmaps of RNA-seq normalized counts were generated using pheatmap. GO ontology analysis was performed usisng clusterprofiler. Gene tracks were generated using the UCSC Genome Browser Custom Tracks feature. Plots of HDAC2 occupancy near transcription start sites were generated using deeptools.

#### Western blotting

To prepare total cell lysate, cells were incubated in ice-cold RIPA buffer containing cOmplete protease inhibitor (Roche) for 10 minutes with rotation. Lysates were then cleared by centrifugation at 11,000 x *g*. Protein concentrations for total or fractionated cell lysates were determined by BCA assay (Fisher Scientific). Equal quantities (20-50 μg) of samples were heated at 95 degrees for 8 minutes in 1X LDS buffer with 1X reducing agent (Thermo Fisher). Samples were then separated by SDS-PAGE and transferred onto an a PVDF membrane (Bio-Rad). Membranes were blocked using 5% BSA in 1X PBS + 0.1% Tween-20 (PBS-T), and subsequently incubated with primary antibody in 5% BSA + PBS-T. Chemiluminescent signal was detected by HRP-conjugated secondary antibodies diluted in 5% BSA + PBS-T, and incubation with ECL western blotting substrate (Pierce). Membranes were stripped using Restore western blot stripping buffer (Pierce) and reprobed as necessary.

#### Northern blotting

20 μg of total RNA was loaded onto a Urea-acrylamide gel, and electrophoresis was carried out at 200V for 2 hours. RNA was transferred onto a nylon membrane (VWR) and UV crosslinked (240 mJ/cm^2^). Prehybridization of the membrane was performed at 42 degrees with UltraHyb-Oligo buffer (Ambion). DNA oligonucleotides complementary to specific RNAs (see supplementary methods) were radiolabeled with [^32^P]ATP using T4 PNK (NEB), and purified using G-50 columns (Fisher Scientific). Blots were incubated with radiolabeled probes overnight at 42 degrees with rotation. The next day, blots were washed once with 2X SSC + 0.1% SDS for 30 minutes, and once with 1X SSC + 0.1% SDS for 30 minutes prior to imaging.

#### Immunofluorescence

MDA-MB-231 cells on 8-well chamber slides (Thermo Fisher) were treated for 30 minutes with 100 nM Mitotracker Deep Red FM in D10F (Thermo Fisher). Cells were then washed twice with chilled 1X PBS, and fixed with 4% paraformaldehyde. Samples were permeabilized on ice for 10 minutes using PBS + 0.5% TritonX-100 and blocked by incubation with 5% goat serum in PBS with 0.1% TritonX-100 (blocking buffer) for 1 h at room temperature. Subsequently, cells were stained with primary antibody diluted in blocking buffer at 4 °C overnight. Slides were then washed three times with PBS + 0.1% TritonX-100, incubated with secondary antibody diluted in blocking buffer for 1 hour at room temperature. Slides were subsequently washed with PBS three times and nuclei were counterstained with DAPI (2.5 μg ml^−1^, Roche) before mounting with Prolong Gold (Thermo Fisher Scientific). Images were acquired using an RS-G4 confocal microscope (Caliber I.D.). Images were analyzed and quantified using ImageJ.

#### tRNA profiling

tRNA profiling was performed as previously described (Goodarzi et al., 2016). Briefly, 300 ng of small RNA (isolated using the microRNA purification kit [Norgen]) was deacylated, biotinylated, and incubated with pooled tRNA probes. Hybridized probes were ligated using T4 DNA ligase and SplintR ligase (NEB), and hybrids were purified using MyOne-C1 Streptavidin Dynabeads (Invitrogen). Ligated probes were eluted by treatment with RNase H and RNase A, PCR amplified, and subjected to high-throughput sequencing. For computational analysis, fastq files were aligned to tRNA probe sequences using bowtie2, and reads were further sorted, indexed, and counts were generated with samtools. Raw counts were imported into R and normalized with DESeq2.

#### Fluorescence *in situ* hybridization

Cells plated into 8-well chamber slides were washed, fixed in 4% paraformaldehyde for 10 minutes at room temperature, and permeabilized using 1X PBS + 0.2% TritonX-100 + 5% acetic acid. An acetylation stock solution was prepared (0.185 g triethanolamine, .224 mL 0.1M NaOH in 10 mL RNase-free water), and 25 µL of acetic anhydride was added immediately prior to use. Samples were incubated in acetylation solution for 10 minutes at room temperature, washed in PBS, and pre-hybridized for 1 hour at 65 degrees in 1X ISH buffer (Qiagen). Digoxigenin-labeled LNA probes against mito-tRNA Asn and nuclear-encoded tRNA Asn designed by Qiagen (sequences in supplementary methods) were boiled for 2 minutes, and then immediately chilled on ice. LNAs were resuspended in 1X ISH buffer at a final concentration of 10-40 nM, and applied to slides overnight at 55 degrees. Slides were subsequently washed at room temperature with 4X SSC + 50% formamide, 2X SSC + 50% formamide, 1X SSC + 50% formamide, and 0.5X SSC + 50 % formamide for 20 minutes each. Samples were then washed once with 100 mM Tris-HCl pH 7.4 + 150 mM NaCl (TN buffer) for 5 minutes, and blocked with TNB buffer (TN buffer + Blocking Reagent [Roche]) for 1 hour at room temperature. Anti-DIG-POD antibody (Roche) was diluted 1:500 in TNB buffer, and incubated on slides overnight at 4 degrees. Slides were then washed three times with TNT buffer (TN buffer + 0.05% Tween-20) and incubated with FITC-tyramide diluted 1:100 in Amplification reagent (PerkinElmer) for 10 minutes at room temperature. Peroxidase activity was quenched with 3% H2O2 at room temperature for 1 hour, slides were counterstain with DAPI, and mounted with Prolong Gold reagent.

#### ρ0 cell generation and ethidium bromide treatment

MDA-MB-231 cells were incubated with 50 ng/mL of ethidium bromide and passaged for ∼6 weeks in D10F supplemented with uridine (50 mg/mL, Sigma). Upon confirmation of loss of mitochondrial transcriptional activity by qPCR and Northern blotting, ethidium bromide was omitted from the media. In parallel, wild type MDA-MB-231 cells were maintained along with the ρ0 cells, to act as a passage-matched control. For acute depletion of mitochondrial transcripts, MDA-MB-231 cells were treated with 200 ng/mL of ethidium bromide for 24 hours, and RNA was isolated for Nothern blot analysis.

#### Cloning

For tRNA overexpression, four tandem repeats of a mito-tRNA Asn or Trp construct were synthesized as previously described ^56^ and cloned into the pUC-GW vector (Genewiz). The NARS2 ORF (Genecopoeia) was used as a template for PCR, and restriction enzyme sites for NotI and EcoRV were introduced for cloning into the pcDNA4 myc/His backbone (Thermo Fisher). For NARS2-Apex2 cloning, restriction enzyme sites for NotI and BamHI were introduced in the NARS2 template for cloning into the pcDNA3-V5-Apex2 backbone_46._

#### Proximity ligation assay

Samples were prepared using the DuoLink PLA reagents (Sigma) following the manufacturer’s protocol. Briefly, cells were fixed and permeabilized, and blocking was performed using DuoLink Blocking solution for 1 hour at 37 degrees. Samples were incubated in primary antibodies diluted 1:5000 in DuoLink Antibody Diluent overnight at 4 degrees. Slides were washed twice with Wash Buffer A, and PLUS and MINUS probes were diluted 1:5 in Antibody Diluent and applied to samples for 1 hour at 37 degrees. Ligation, amplification, and washing was then performed as per the manufacturer’s protocol, and samples were counterstained with DAPI and mounted using DuoLink In Situ Mounting Media.

#### Oxygen consumption and extracellular acidification rate measurements

Oxygen consumption and Extracellular acidification rates (OCR and ECAR respectively) were measured in MDA-MB-231 cells using the XF96 Extracellular Flux analyzer (Seahorse Bioscience). Briefly, 2 x 10^4^ cells per well were seeded in XF96 cell culture multi-well plates in growth medium and incubated overnight. XF96 cartridges were hydrated overnight in deionized water at 37°C in a non-CO2 incubator. Prior to OCR measurements, the growth medium of cells was exchanged with XF medium supplemented with glucose, glutamine and pyruvate and incubated at 37°C in a non-CO2 incubator for 1 hr. Inhibitors were diluted to appropriate concentrations in XF medium and loaded into corresponding microwells in the XF96 cartridge plate. Sensor cartridges were equilibrated in XF calibrant medium and calibrated in the Flux analyzer. XF96 cell culture plate was loaded into the Flux analyzer and OCR and ECAR were measured. Data were normalized to cell numbers determined by counting NucBlue–stained nuclei using the ImageXpress micro imaging system.

#### Y-shaped Adapter-ligated Mature tRNA (YAMAT) qPCR

We adapted the YAMAT sequencing protocol ^86^ to quantify the relative abundance of mature mito-tRNAs where indicated. Briefly, we enriched for small RNAs from total RNA samples. For each condition, 500ng of small RNAs was deacylated in 20mM Tris-HCl (pH 9.0) at 37°C for 40–60 minutes and demethylated using rtStar tRNA-optimized First strand cDNA synthesis kit following manufacturer’s protocol with intervening RNA cleaning steps (Zymo RNA Clean and Concentration Kit). For the annealing reaction, 1µL of adapter duplexes made up of IDT-synthesized 40µM DNA adapter containing Illumina 3’-adapter sequences (Y-3’-AD) and 10µM each of four DNA/RNA hybrid adapters containing Illumina 5’-adapter sequences (Y-5’-AD) with 3’-terminal A, G, C, or U was incubated with 8µL of deacylated and demethylated small RNAs at 90°C for 2 minutes. The adapter sequences were as follows:

Y-3’-AD: 5’-5phos/GTATCCAGTTGGAATTCTCGGGTGCCAAGG/3ddC-3’ Y-5’-AD-A: 5’-GTTCAGAGTTCTACAGTCCGACGATCACTGGATACTGga-3’ Y-5’-AD-G: 5’-GTTCAGAGTTCTACAGTCCGACGATCACTGGATACTGgg-3’ Y-5’-AD-C: 5’-GTTCAGAGTTCTACAGTCCGACGATCACTGGATACTGgc-3’ Y-5’-AD-U: 5’-GTTCAGAGTTCTACAGTCCGACGATCACTGGATACTGgu-3’

While the RNA-Adapter mixture was incubated at 37°C, 1µL of pre-warmed (37°C)10x annealing buffer (50 mM Tris HCl pH 8, 100 mM MgCl_2_, 5 mM EDTA) was added to the reaction and incubated at 37°C for a further 15 minutes. To ligate the annealed adapters to mature tRNAs, 10µL of ligation reaction mix (1µL of 10x RNA ligase buffer, 0.1µL Rnl2, 0.4µL of 10mM ATP, 0.1µL of 100mM DTT and 8.4µL nuclease-free water) was added to the RNA-adapter reaction mixture and incubated at 37°C for 1 hour followed by 4°C overnight incubation. cDNA was synthesized from adapter-ligated tRNAs using the First strand cDNA synthesis Kit (ThermoFisher) using the following primer for reverse transcription: 5’-GCCTTGGCACCCGAGAATTCCA-3’. Fragments larger than 100bps were purified using Ampure XP beads and 20µL of purified tRNAs was diluted 1:15 in nuclease-free water and relative mito-tRNA abundance in each sample was determined by qPCR as described above.

#### APEX2 Proteomics and Data Analysis

About 70-80% confluent HEK293T cells were transiently transfected with APEX2 fusion constructs using Lipofectamine 3000 and incubated for 48 hours. Transfected cells were loaded with 500uM Biotin phenol or DMSO control and incubated for 30 min at 37°C. H2O2 was added to the cell culture media to a final concentration of 1 mM to induce Apex2-mediated biotinylation for 1 min with gentle swirling. The media was aspirated and the cells were washed three times with quencher solution containing 10 mM sodium ascorbate, 10 mM sodium azide, and 5 mM Trolox in phosphate-buffered saline (PBS). The cells were scraped into ice-cold PBS, transferred to falcon tubes and centrifuged at 3,000 × g for 5 min. To enrich for biotinylated proteins, cell pellets were lysed in RIPA lysis buffer supplemented freshly with Halt protease and phosphatase inhibitor cocktail (Themo Scientific), 10 mM sodium ascorbate, 10 mM sodium azide, and 5 mM Trolox for 10 minutes on ice. The lysates were clarified by centrifugation at 15000 x g at 4°C for 10 minutes. Streptavidin-coated magnetic beads (Pierce) were washed twice in RIPA lysis buffer. 1mL total volume of 10mg protein lysate from each condition and the prepared magnetic beads was rotated overnight at 4°C. The beads were pelleted on a magnetic rack, washed twice with RIPA lysis buffer, once with 1M KCl, once with 0.1 M Na_2_CO_3_, once with 2 M urea in 10 mM Tris-HCl (pH 8.0) and twice with RIPA lysis buffer at 4°C. The beads were dried on the magnetic rack. Proteins coupled to streptavidin-coated magnetic beads were reduced with dithiothreitol (Sigma-Aldrich) and alkylated with iodoacetamide. Proteins were eluted from beads using partial on-bead trypsination (Promega) followed by a second digestion. Samples were micro solid phase extracted (Empore, 3M) ^87^ and analyzed by LC-MS/MS: 50 min analytical gradient (2%B to 38%B, A: 0.1% formic acid, B: 80% acetonitrile, 0.1% formic acid), 12 cm built-in-emitter column, high resolution/high mass accuracy (Orbitrap Fusion LUMOS, Thermo). The data were processed using Proteome Discoverer 1.4 /Mascot as well as MaxQuant ^88^ v.1.6.6.0 and queried against UniProt’s Human databases concatenated with common contaminants. Perseus ^89^ v.1.6.10.50 were used for data analysis. Matched proteins were filtered for possible contaminants. Proteome Discoverer 1.4 protein abundance (average-of the-3-most abundant-peptides for a matched protein) were log2 transformed and further filtered requiring that a protein signal must be present in minimum 2/3 of the replicates for at least one condition. Missing values were imputed (width: 0.3, down shift 1.8) followed by a quartile-based width adjustment normalization.

#### Steady state polar metabolite profiling

For steady state polar metabolite profiling, polar metabolites were extracted as described previously ^90^. Briefly, 3.5×10^5^ MDA-MB-231 cells were cultured as triplicates or quadruplicates and transfected as described above. 48 hours post-transfection, the cells were washed three times with 1 ml of cold 0.9% NaCl and scraped into 500µL of cold 80%v/v LC-MS grade methanol containing heavy amino acid internal standards (Cambridge Isotope Laboratories). After 10 minutes of vortexing at 4°C, samples were centrifuged at 16000xg for 10 minutes to remove cell debris. The samples were nitrogen-dried and stored at −80 °C until analysis by LC-MS. The dried pellets were resuspended in 60 μl of 50% acetonitrile, vortexed vigorously for 10 seconds followed by centrifugation at 16,000 RCF at 4°C for 30 minutes. The supernatant (5 μl) was injected onto a ZIC-pHILIC 150 × 2.1 mm (5-μm particle size) column (EMD Millipore) connected to a Thermo Vanquish ultrahigh-pressure liquid chromatography (UPLC) system and a Q Exactive benchtop orbitrap mass spectrometer equipped with a heated electrospray ionization (HESI) probe. The samples were analyzed via LC-MS/MS in a randomized sequence. Integration of extracted ion chromatograms was performed using Skyline Daily ^91^ (v 21.1) with the maximum mass and retention time tolerance set to 2ppm and 12sec, respectively, referencing an in-house library of chemical standards. Metabolite levels (peak areas) were normalized to the median metabolite signal within each sample. The normalized area-under-the-curve (AUC) was used for relative quantification. Statistical and metabolite set enrichment analyses were performed using MetaboAnalyst ^92^ v 5.0.

#### Quantification and statistical analysis

Unless otherwise noted all data are expressed as mean ± standard error of the mean. Groups were compared using tests for significance as indicated in the figure legends and the text. A significant difference was concluded at P < 0.05. Where indicated in figures: *P < 0.05, **P < 0.01, ***P < 0.001, and ****P < 0.0001.

## ACKNOWLEDGEMENTS

We thank members of our laboratory for comments on the manuscript. We are grateful for assistance by Rockefeller University resource centers: Connie Zhao and staff at the genomics resource center for assistance with sequencing experiments, and Vaughn Francis and other veterinary staff of the Comparative Bioscience Center for animal husbandry and care. We acknowledge the outstanding technical contributions of Ophelia Lee for molecular cloning of NARS2 constructs, and Greta Ghita for contributions to sequencing analyses. This work was supported by an R35CA274446 award to S.F.T and the Black Family Center for Human Metastasis Research. The Rockefeller University Proteomics Resource Center acknowledges funding from the Leona M. and Harry B. Helmsley Charitable Trust and Sohn Conferences Foundation for mass spectrometer instrumentation.

## AUTHOR CONTRIBUTIONS

S.F.T. conceived the study and supervised all research. C.R., K.F.Y., and S.F.T. designed all experiments. C.R., K.F.Y., M.L.D., H.A., and H.M. conducted experiments. C.R. analyzed sequencing data. C.R. and S.F.T. wrote the manuscript with input from co-authors.

## Supplementary Figures and Figure Legends

**Figure 1.**
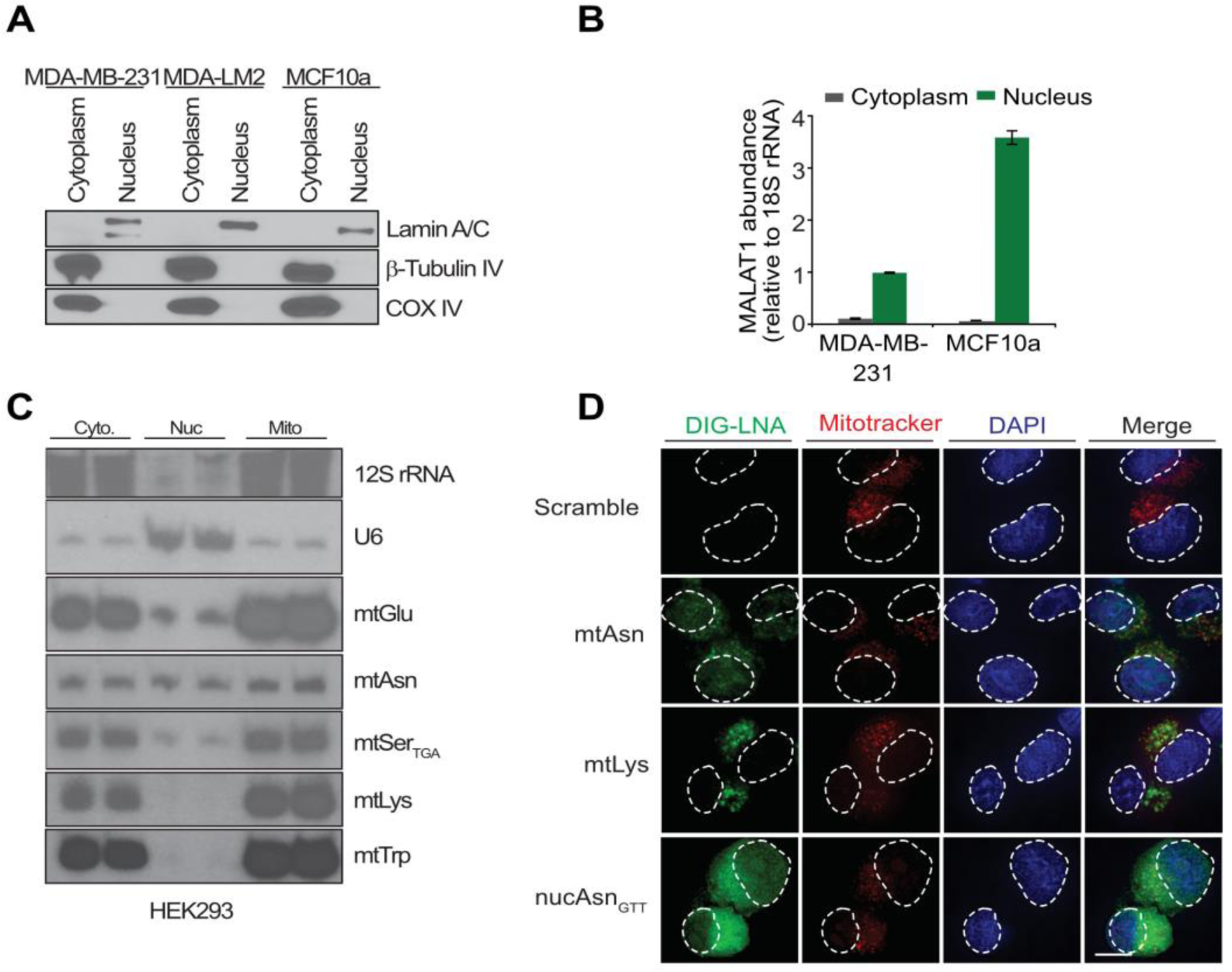
Fractionation and validation of cell lines to detect nuclear-localized mito-tRNA. A. Subcellular fractionation and immunoblotting was performed on indicated cells for the specified proteins. B. RT-qPCR for the nuclear MALAT1 long, non-coding RNA from purified nuclear or cytoplasmic/mitochondrial total RNA. C. Northern blot analysis of cytoplasmic/mitochondrial and nuclear fractions of total RNA from HEK293T cells to detect abundance of labeled RNAs D. RNA FISH of mito-tRNA (Asn, Lys) and nuclear-encoded tRNA Asn_GTT_ localization in MDA-MB-231 cells. DAPI and Mitotracker Deep Red visualize the nuclei and mitochondria of cells, respectively. Dotted outlines highlight nuclear position, scale bar = 10 μm.

**Figure 2.**
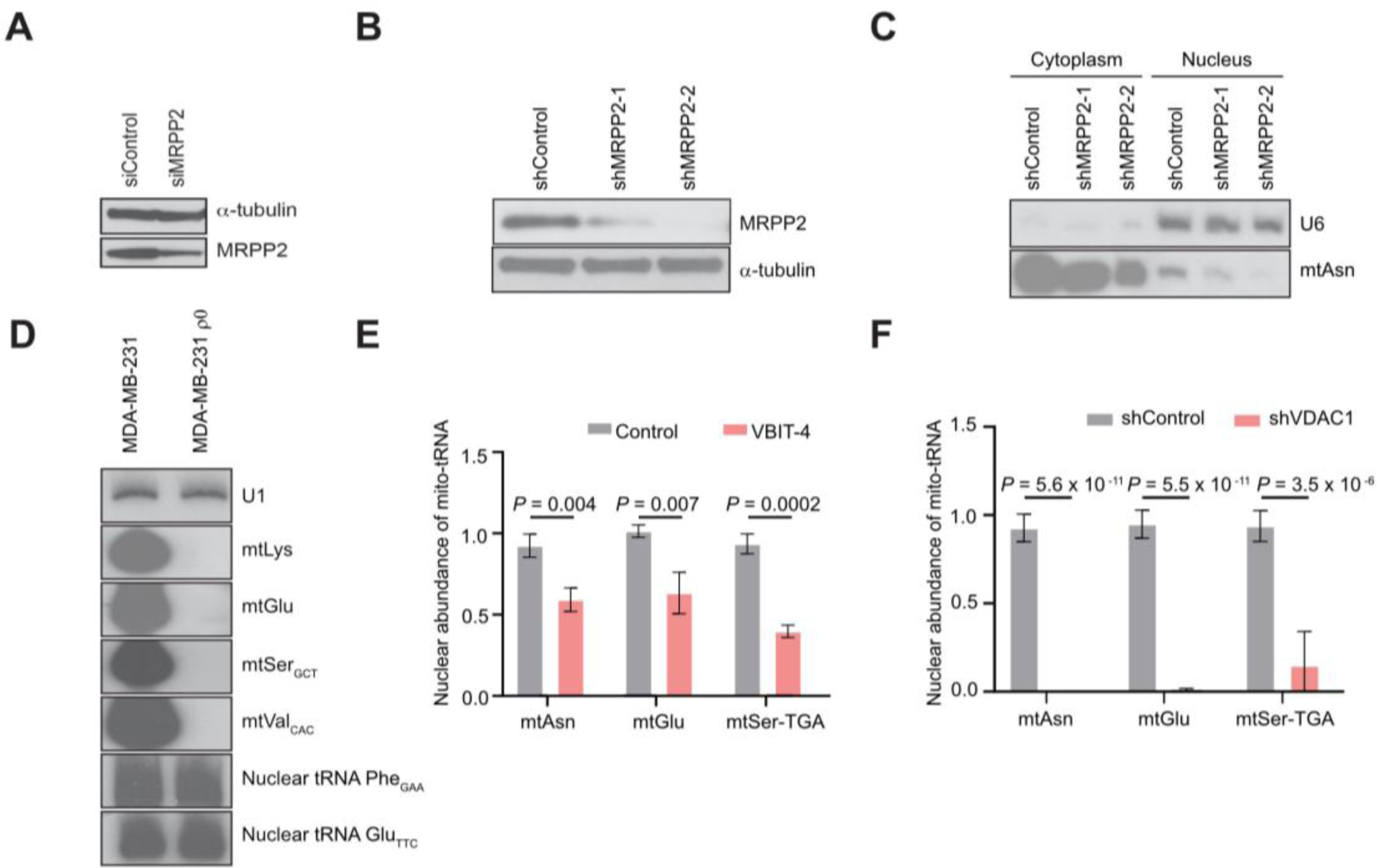
Nuclear mito-tRNA abundance depends on mitochondrial tRNA transcription, processing and the VDAC1 pore. A. siRNA-mediated depletion of MRPP2 in HEK293 cells. Total protein lysate was analyzed from HEK293 cells transfected with scramble control siRNA, or one targeting MRPP2 by immunoblotting to determine degree of MRPP2 depletion. B. MDA-MB-231 infected with lentivirus encoding shRNA cassettes against MRPP2 were lysed, and protein content was assessed for MRPP2 reductions by immunoblotting. C. Total RNA derived from the cytosolic/mitochondrial or nuclear fractions of MDA-MB-231 cells expressing non-targeting, or MRPP2-directed shRNAs was subjected to Northern blot analysis for the indicated RNAs. D. Total RNA from MDA-MB-231 or MDA-ρ0 cells was subjected to Northern blot analysis for the labeled RNAs. E. Nuclear abundance of indicated mito-tRNAs measured by YAMAT-qPCR in VBIT-4-treated cells, *n* = 3. F. Nuclear abundance of indicated mito-tRNAs measured by YAMAT-qPCR in shVDAC2 cells, *n* = 6. P-values for (E) and (F) calculated using two-sided Student’s t-test.

**Figure 3.**
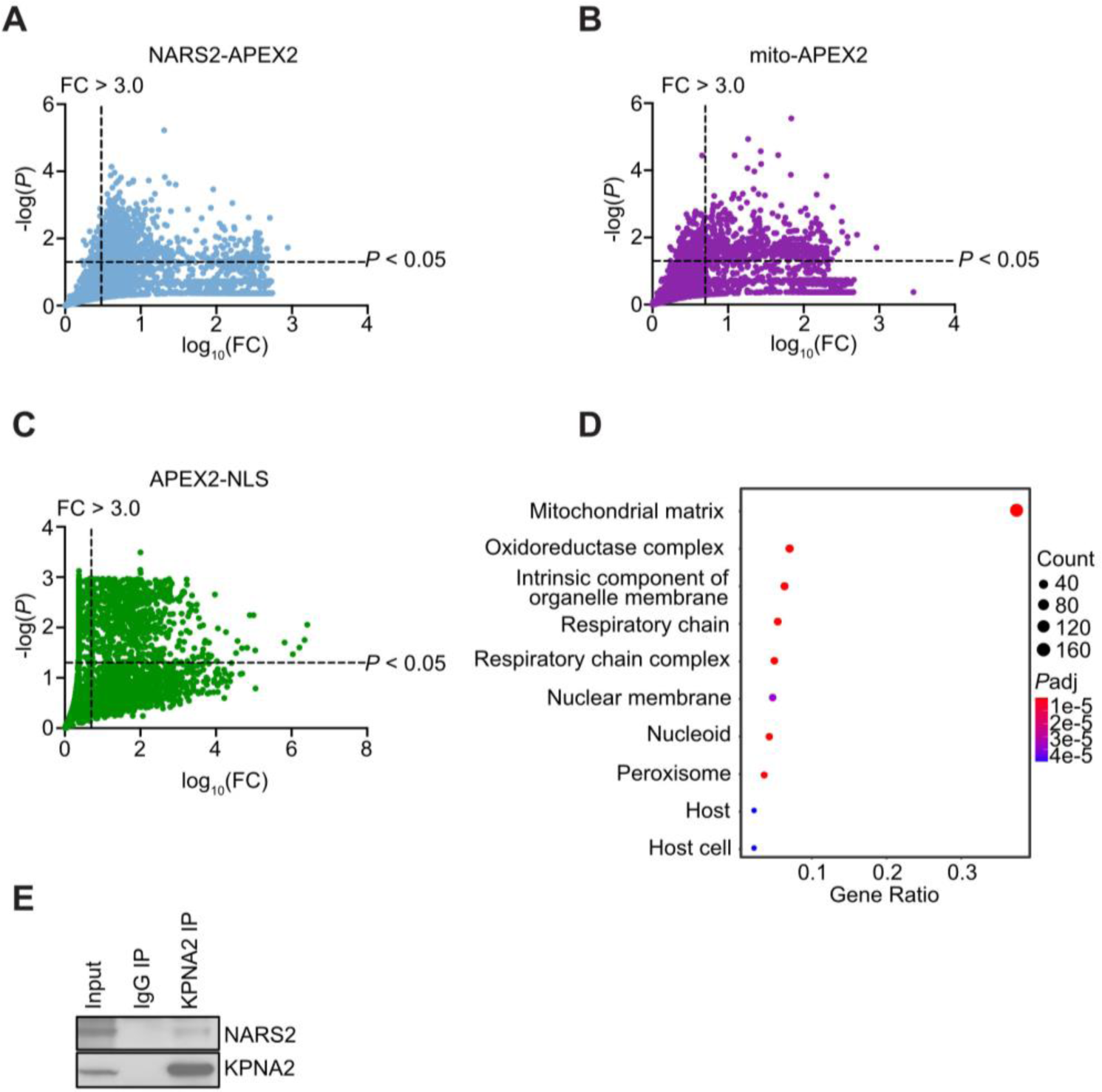
APEX2 proximity biotinylation and mass spectrometry. A, B, C. Plots of proteins identified by mass spectrometry from (**A**) NARS2-APEX2 (**B**) Mito-APEX2 and (**C**) NLS-APEX2 experiments. Dotted lines represent cutoffs for indicated fold changes and p-values. D. GO cell compartment classification of top 500 (logFC ranked, p-value <0.05) proteins identified from NARS2-APEX2 experiments. E. KPNA2 immunoprecipitation was performed, and immunoblotting carried out for the labeled factors.

**Figure 4.**
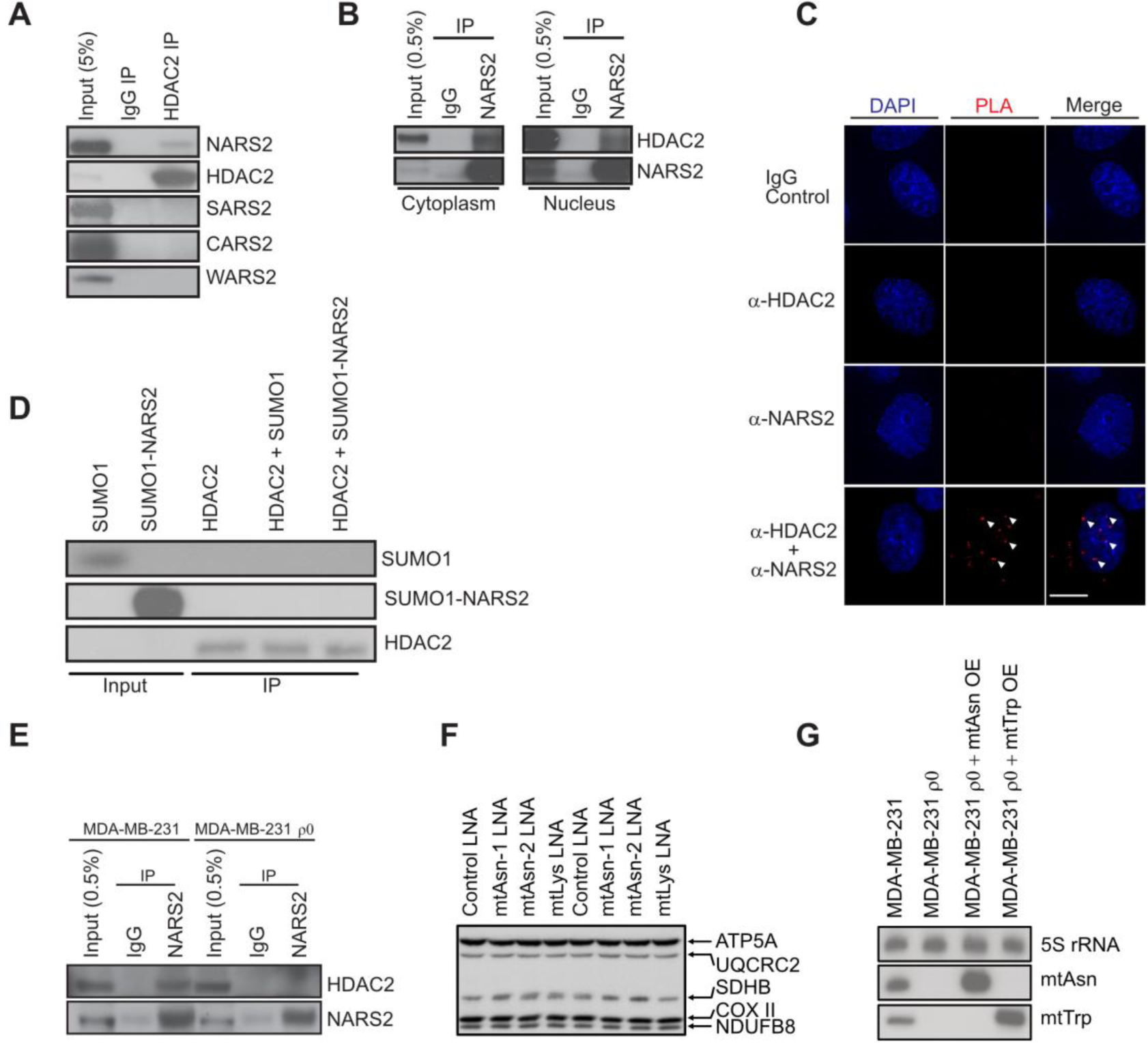
Immunoprecipitation assays to detect NARS2-HDAC2 binding. A. HDAC2 immunoprecipitates were subjected to immunoblot analysis for the labeled proteins. B. Nuclear and cytosolic/mitochondrial MDA-MB-231 cell lysates were used to perform NARS2 immunoprecipitations followed by immunoblotting for the indicated factors. C. Images from proximity ligation assays, where fixed cells were incubated with the indicated antibodies. D. Recombinant HDAC2 and SUMO-tagged NARS2 were incubated *in vitro*, and HDAC2 immunoprecipitations were performed. Immunoblotting for the indicated factors was then carried out. E. NARS2 immunoprecipitations were performed from MDA-MB-231 and MDA-MB-231-ρ0 lysates, and immunoblotting of indicated proteins was carried out. F. Northern blot analysis of MDA-MB-231, MDA-MB-231 ρ0 cells, and MDA-MB-231 ρ0 cells with rescued overexpression of mtAsn and mtTrp.

**Figure 5.**
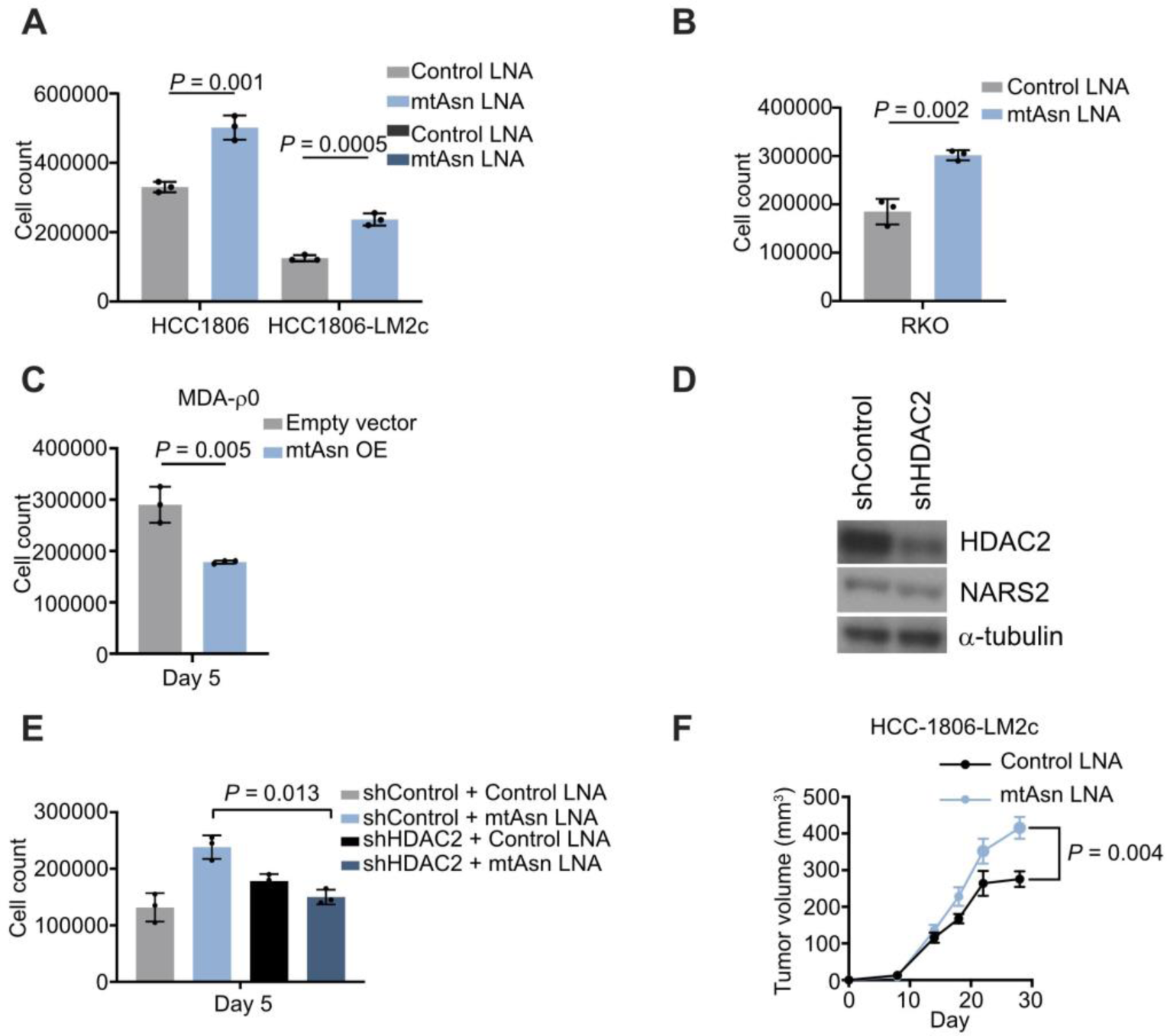
mtAsn inhibition represses target gene expression. A. HCC1806 and HCC1806-LM2c cells transfected with control and mtAsn LNAs were seeded in 6-well plates, and counted 5 days later. n = 3, p-values determined by two-sided Student’s t-test. B. Proliferation assays were performed as in (**A**), but using RKO colon cancer cells. n = 3, p-values determined by two-sided Student’s t-test. C. *In vitro* proliferation assay upon overexpression of mtAsn, n = 3, p-values calculated using two-sided Student’s t-test. D. Immunoblot of lysates derived from cells expressing non-targeting or HDAC2-targeting shRNAs for the indicated proteins. E. Proliferation assays in shControl or shHDAC2 cells upon transfection of the indicated LNAs, n = 3, p-values determined by two-sided Student’s t-test. F. 50,000 HCC-1806-LM2c cells transfected with indicated LNAs were transplanted into the mammary fat pads of mice, and primary tumor growth measured. n = 7 per group, p-value calculated by two-sided Student’s t-test.

**Figure 6.**
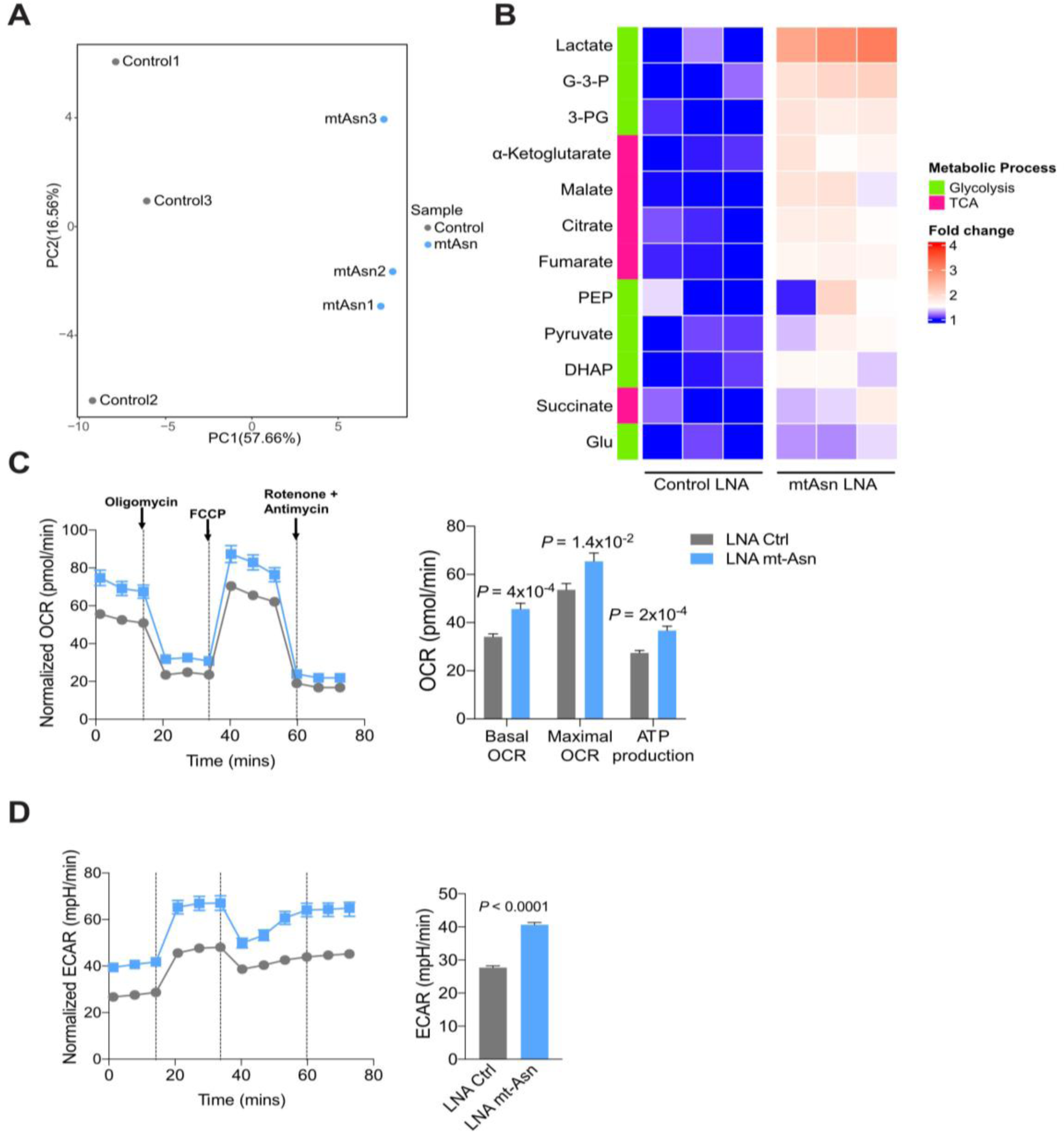
Metabolic control via mtAsn. A. PCA analysis for steady-state metabolite abundance in MDA-MB-231 cells transfected with control or mtAsn LNAs. B. Heat map representation of total metabolite abundance for glycolytic and TCA cycle intermediates in MDA-MB-231 cells transfected with control or mtAsn LNAs. C. Oxygen consumption rate (OCR) in MDA-MB-231 cells transfected with control or mtAsn LNAs. D. Extracellular acidification rate (ECAR) in MDA-MB-231 cells transfected with control or mtAsn LNAs.

**Figure 7.**
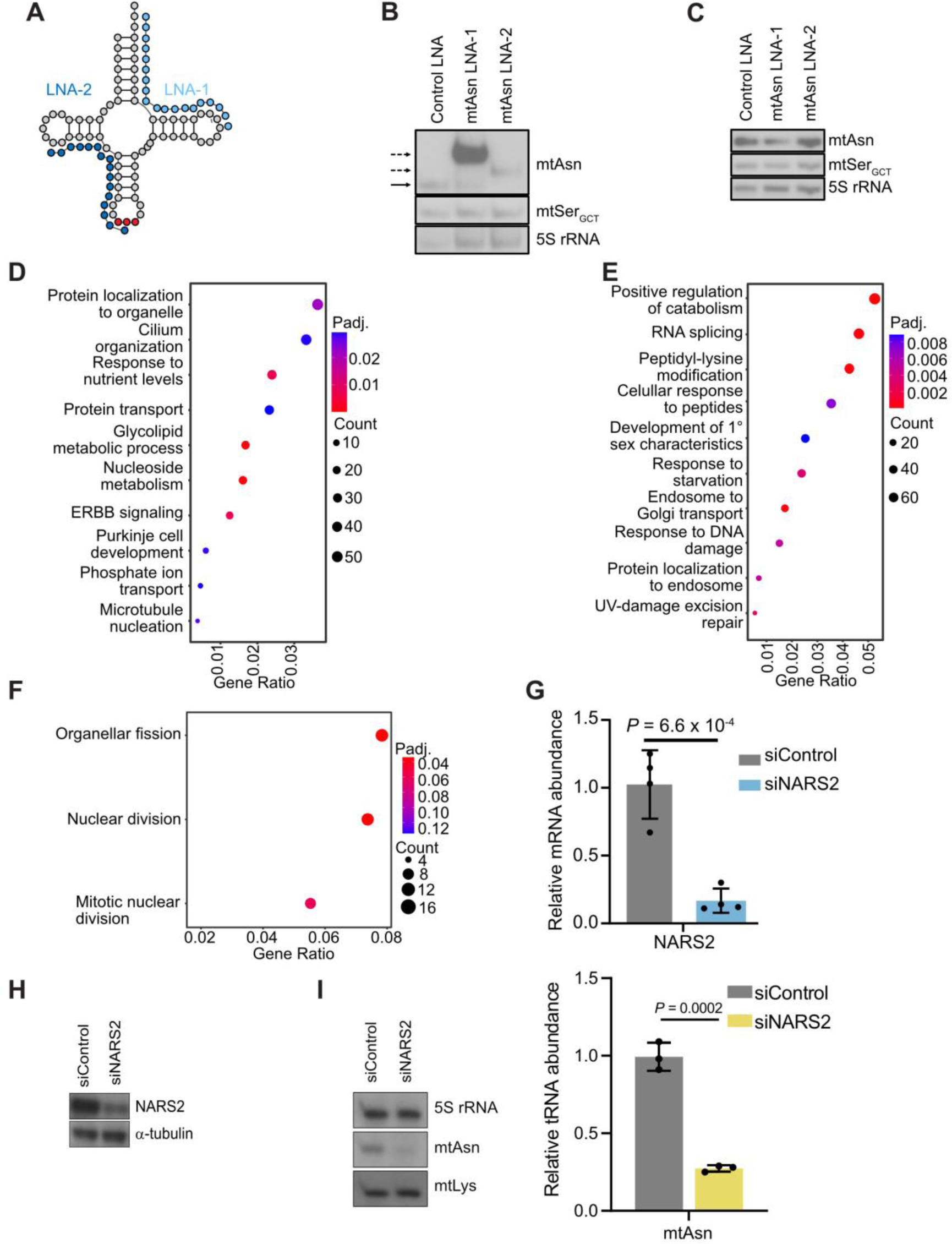
mtAsn inhibition regulates target gene expression. A. Schematic representation of mtAsn and targeting LNAs. B. Northern blotting of total RNA from MDA-MB-231 cells transfected with non-targeting LNA, mtAsn LNA-1, or mtAsn LNA-2 under native conditions for the indicated RNAs. Solid and dashed arrows indicate positions of mtAsn alone or band-shifted, LNA-bound complexes, respectively. C. Northern blotting of total RNA from MDA-MB-231 cells transfected with non-targeting LNA, mtAsn LNA-1, or mtAsn LNA-2 under denaturing conditions for the indicated RNAs. D. GO pathway enrichment of genes commonly upregulated by mtAsn and mtLys inhibition. E. GO pathway enrichment of genes commonly downregulated by mtAsn and mtLys inhibition. F. GO pathway enrichment of upregulated mtAsn-specific genes. G. RT-qPCR analysis of RNA isolated from MDA-MB-231 cells transfected with negative control or NARS2-targeting siRNAs. n = 4, p-values calculated using two-sided Student’s t-test. H. Immunoblot of lysates derived from MDA-MB-231 cells transfected with siRNAs targeting NARS2 or a non-targeting negative control. I. Northern blot (left) and quantification (right) of indicated RNAs derived from cells transfected with negative control and NARS2 siRNAs.

**Figure 8.**
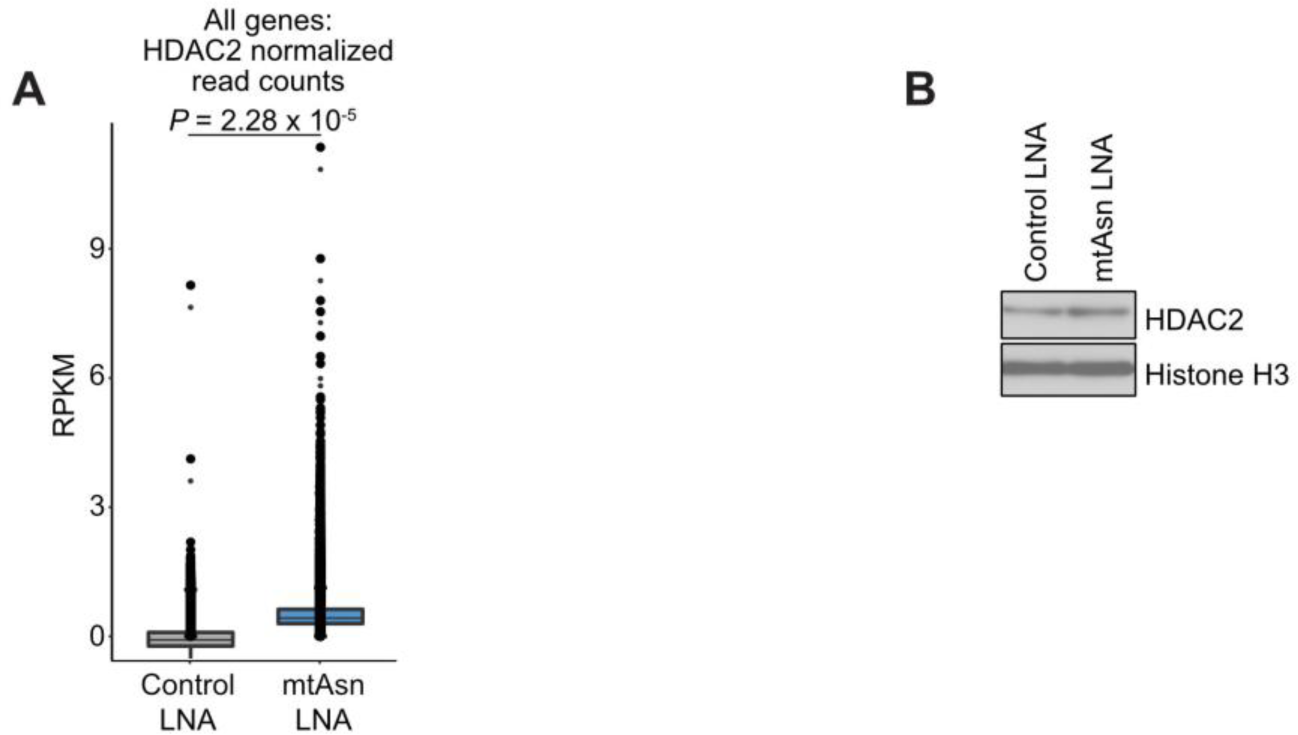
Metabolic control via mtAsn. A. Normalized count measurements at HDAC2 differentially bound regions. Normalization was performed using RPKM-based calculation, and p-value was determined by two-sided Student’s t-test. B. Immunoblot of chromatin-associated HDAC2 in MDA-MB-231 cells upon transfection with control LNA or mtAsn LNA.

**Table S1.**
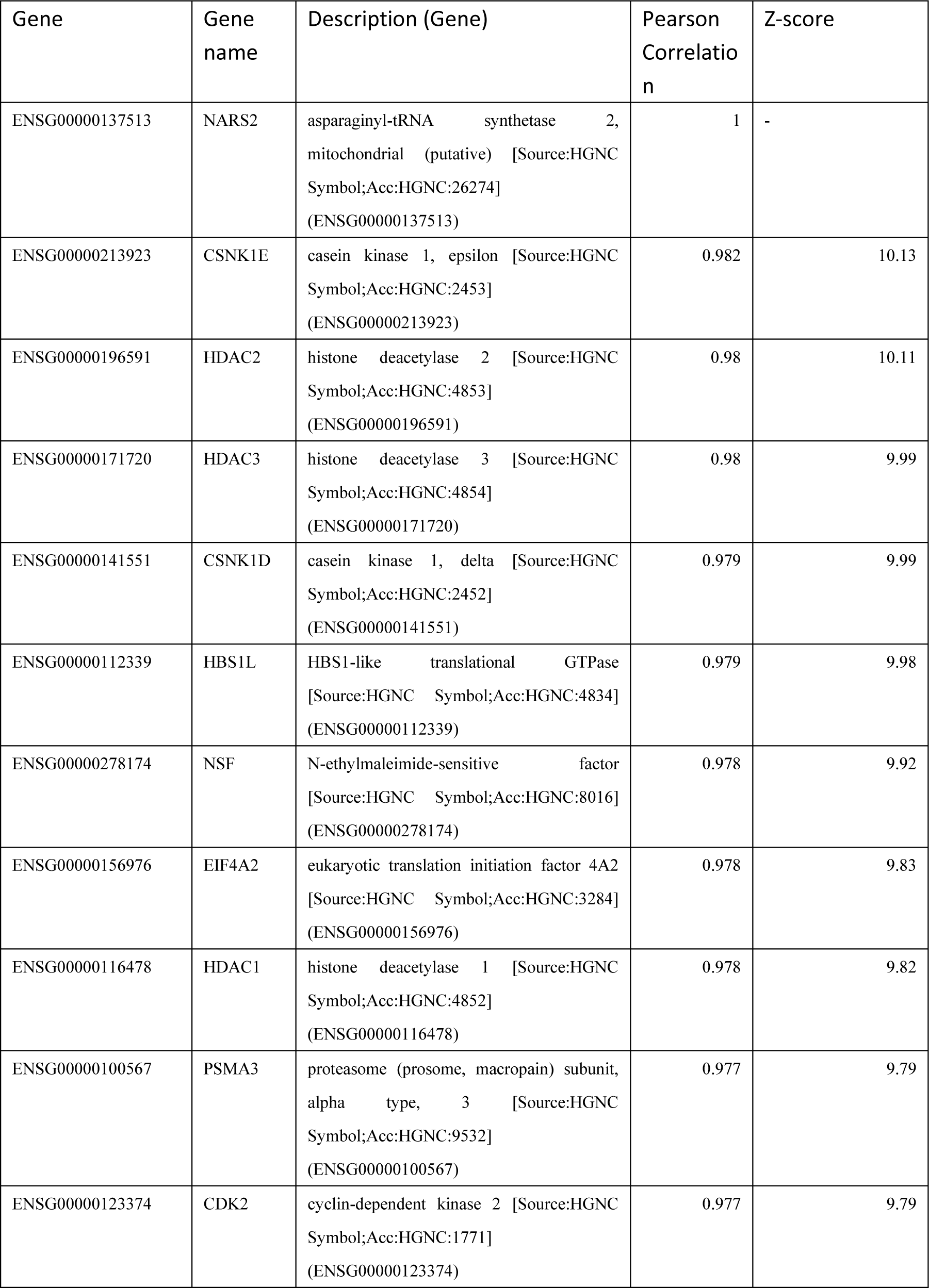
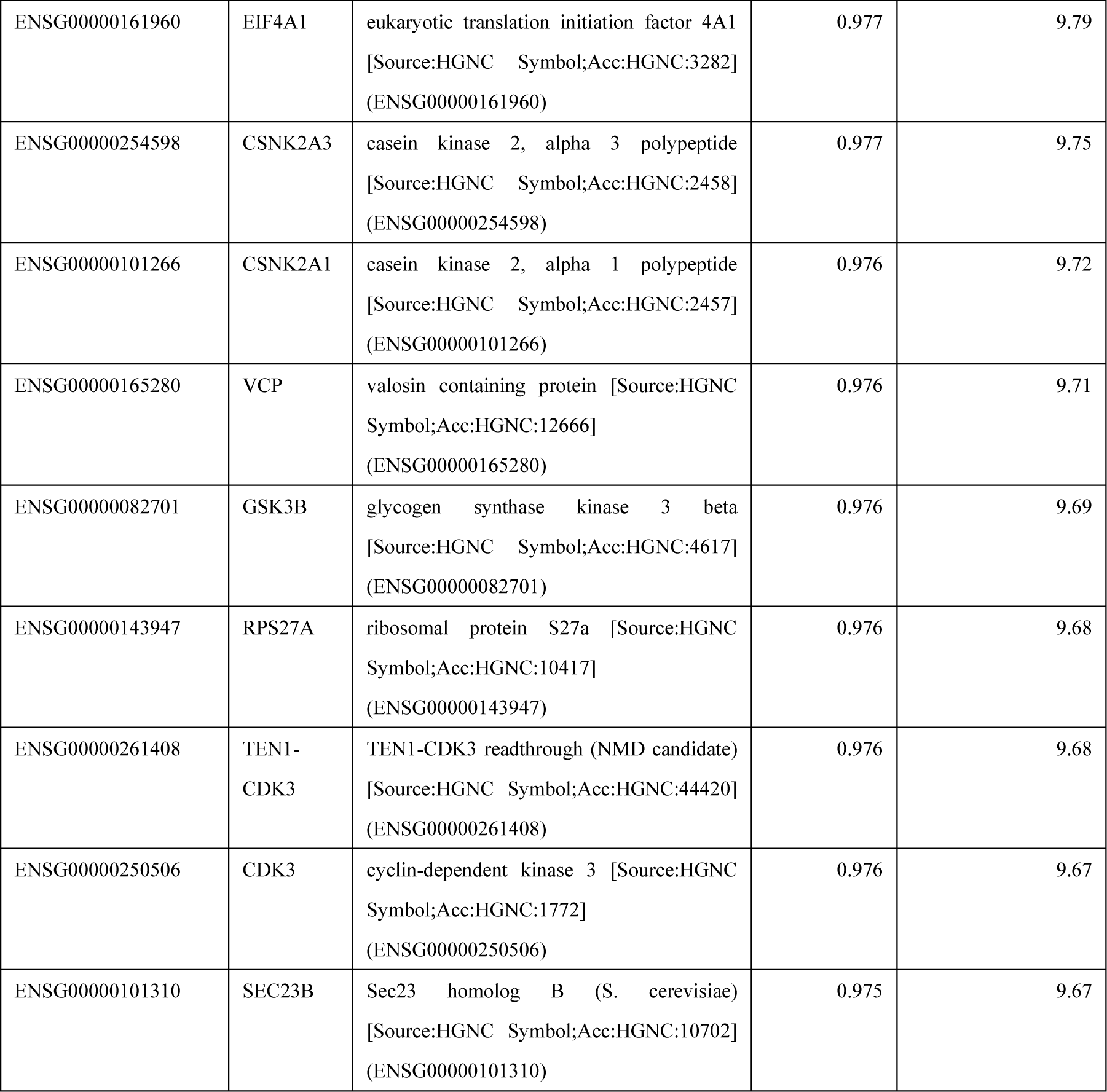
Genes with co-evolving protein sequences as calculated using Phylogene.

